# Tumour-derived gliogenesis sustains dedifferentiation-dependent tumour growth in the *Drosophila* CNS

**DOI:** 10.1101/2024.12.23.630170

**Authors:** Edel Alvarez-Ochoa, Qian Dong, Hannah Truong, Louise Y Cheng

## Abstract

Fate-restricted cells can acquire stem cell-like properties through dedifferentiation, enabling them to gain the plasticity required for differentiation into multiple lineages. Tumour plasticity is prominently observed in brain cancers, where transient cell state changes are linked to resistance to conventional therapies. In this study, we demonstrate that a sub-population of dedifferentiated tumour neural stem cells (NSCs) in *Drosophila*, induced by the knockdown of *prospero* (*pros*), can generate its own glial niche. Temporal patterning, known to influence oncogenic competence and tumour malignancy, plays a key role in this process. Specifically, we show that *de novo* gliogenesis occurs in the more differentiated Syncrip+ (Syp^+^) NSC population. Modulating Syp levels alters the size of the glial niche, subsequently affecting tumour size. Furthermore, the tumour-associated glial niche expands through cell division and fails to cease proliferation on time due to dysregulated ecdysone signalling, contributing to niche expansion. Our findings reveal that tumours arising via dedifferentiation establish their own supportive glial microenvironment, which sustains tumour growth.

## Introduction

Neural stem cells (NSCs) are surrounded by a niche made up of cellular and extracellular components which communicate with NSCs to regulate their behaviour. The neurogenic niche of NSCs can secrete growth signalling factors to promote or inhibit NSC proliferation, or spatially anchor NSCs to their native microenvironment (Li & Guo, 2021; Morante-Redolat & Porlan, 2019). Dedifferentiation is the process by which mature cells revert to a stem cell-like state, reactivating genes associated with multipotency and leading to tumour formation. This dysregulation underlies the developmental origin of many childhood brain cancers (Friedmann-Morvinski *et al*, 2012). Tumour progression has also been associated with changes in the neurogenic niche. For instance, glioma cancer stem cells induce astrocyte reactivation which in turn promotes tumour growth and metastasis (Oushy *et al*, 2018; Zhang *et al*, 2020). Despite these insights, it remains unclear how brain tumours promote changes in the niche to generate an adaptive microenvironment that supports tumour growth, and whether tumour plasticity plays a role in this process.

The cellular dynamics and molecular pathways regulating NSC behaviour have been extensively studied in the *Drosophila* central nervous system (CNS) (Homem & Knoblich, 2012). *Drosophila* NSCs, called neuroblasts (NBs), undergo self-renewal and differentiation during development to generate the neurons and glial cells that constitute the adult brain. Upon asymmetric division, the majority of the NBs located in the central brain (CB) and the ventral nerve cord (VNC) (type I NBs) self-renew and give rise to an intermediate progenitor, called ganglion mother cell (GMC), that divides once to generate two post-mitotic neurons or glial cells (Homem & Knoblich, 2012). In contrast, a small number of NBs in the CB (type II NBs) generate intermediate neural progenitors (INPs) that produce a few GMCs, and a large number of neurons, enabling further expansion of the lineage (Figure 1A) (Homem & Knoblich, 2012). Regulators involved in the differentiation of GMCs and INPs, or the maintenance of neuronal differentiation, are required for restraining the NB proliferative potential and preserving the invariant NB numbers in the brain. The inactivation of the transcription factor Prospero (Pros) in the GMCs (Betschinger *et al*, 2006; Choksi *et al*, 2006), of Nervous fingers-1 (Nerfin-1) (Froldi *et al*, 2015, 2019; Veen *et al*, 2023; Vissers *et al*, 2018), Longitudinals lacking (Lola) (Southall *et al*, 2014) and Midlife crisis (Mdlc) (Carney *et al*, 2013) in the neurons, and of the NHL translation repressor Brain tumour (Brat) in the INPs (Betschinger *et al*, 2006), results in their dedifferentiation into additional NBs. Similarly, disruption to the membrane-targeted polarity protein atypical protein kinase C (aPKC) in the NBs can impair their asymmetric division, resulting in ectopic NBs (Lee *et al*, 2006). These defects can cause an exponential expansion of the stem cell pool, giving rise to brain tumours that exhibit unrestrained proliferative potential and fail to obey signals to undergo terminal differentiation (Caussinus & Gonzalez, 2005).

**Figure 1.**
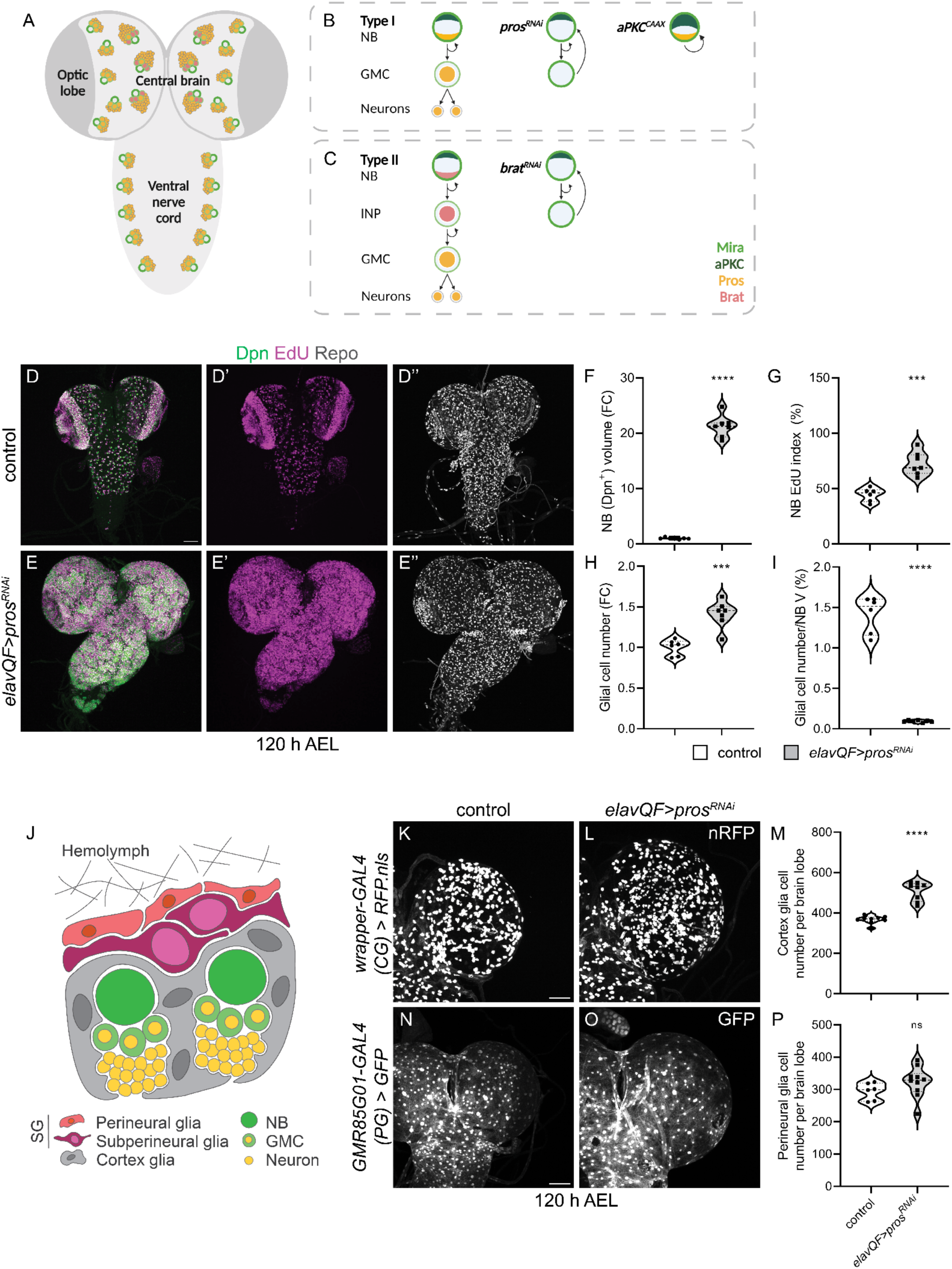
Neuroblast (NB)-derived tumours result in the expansion of the cortex glia (CG) niche. A. Schematic representation of the *Drosophila* larval brain, composed of two optic lobes (OLs), the central brain (CB), and the ventral nerve cord (VNC). B. Schematic representation of type I NB divisions. Type I NBs express cortical Miranda (Mira), apical atypical protein kinase C (aPKC) and basal Prospero (Pros) (left panel). Upon asymmetric division, the NB self-renews and generates a ganglion mother cell (GMC), which inherits Pros. Pros translocation into the nucleus promotes GMC differentiation into two post-mitotic neurons. Expression of *pros^RNAi^* causes GMC to NB reversion (middle panel). Upon hyperactivation of aPKC, NBs fail to differentiate, resulting in the expansion of the NB pool (right panel). C. Schematic representation of type II NB divisions. Type II NBs, which are exclusively located on the dorsal side of the CB, express basal Brain tumour (Brat). Upon asymmetric division, the NB self-renews and generates an intermediate neural progenitor (INP). INPs divide several times while generating GMCs, which express Pros and produce terminally differentiated neurons or glial cells. Expression of an RNAi against Brat causes INP to NB reversion, causing an expansion of the NB pool. D-E’’. Representative images (maximum projections) showing that RNAi-mediated *pros* knockdown in pan-NB lineages via *elav-QF* results in brain overgrowth, expanded NB lineage size (NBs marked by Dpn), increased proliferation (EdU) and increased glial cell number (Repo), at 120 hours after egg laying (h AEL). Scale bar= 50 μm. F. Quantification of Dpn^+^ NB volume in D and E, performed using an unpaired t-test. Data normalised to controls. Control: n= 6, m= 1 ± 0.07. *pros^RNAi^*: n= 7, m= 21.24 ± 0.75. ****p < 0.0001. G. Quantification of NB EdU index in D and E, performed using an unpaired t-test. Control: n= 6, m= 44.39 ± 2.48. *pros^RNAi^*: n= 7, m= 72.54 ± 3.93. ***p < 0.001. H. Quantification of Repo^+^ glial number in D and E, performed using an unpaired t-test. Control: n= 6, m= 1 ± 0.04. *pros^RNAi^*: n= 7, m= 1.41 ± 0.06. ***p < 0.001. I. Quantification of the percentage of Repo^+^ cell number/Dpn^+^ volume in D and E, performed using an unpaired t-test. Control: n= 6, m= 1.42 ± 0.09, *pros^RNAi^*: n= 7, m= 0.09 ± 0.01. ****p < 0.0001. J. Schematic illustrating the glial niche surrounding neural lineages, depicting the surface glia (SG), composed of perineurial glia (pink) and subperineurial glia (purple), the cortex glia (grey), NBs (green), GMCs (green/yellow) and neurons (yellow). K-L. Representative images (maximum projections) showing cortex glia (CG) cells in *pros^RNAi^* brains and control, marked by *wrapper-GAL4>RFP.nls*, at 120 h AEL. Scale bar= 50 μm. M. Quantification of CG cell number in K and L, performed using an unpaired t-test. Control: n= 9, m= 366.7 ± 6.38. *pros^RNAi^*: n= 8, m= 507.8 ± 16.40. ****p < 0.0001. N-O. Representative images (maximum projections) showing perineural glia (PG) cells in control and *pros^RNAi^* brain, marked by *GMR85G01-GAL4>GFP*, at 120 h AEL. Scale bar= 50 μm. P. Quantification of PG cell number in N-O, performed using an unpaired t-test. Control: n= 6, m= 294.3 ± 10.83. *pros^RNAi^*: n= 10, m= 322.7 ± 14.89. ns (not significant) p > 0.05.

*Drosophila* larval NBs reside within a neurogenic niche made up of cortex glial (CG) cells, which individually encase NBs and their newborn progeny within membranous chambers, while forming a network spanning the whole brain (Rujano *et al*, 2022) (Figure 1 J). The CG niche has been implicated in the regulation of NBs’ ability to emerge from quiescence, proliferate, and terminally differentiate (reviewed in (Nguyen & Cheng, 2022)). Its structure changes throughout neural development, implying that its role in NB regulation could evolve according to the needs of the stem cell lineages (Spéder & Brand, 2018). However, how the niche morphology changes during dedifferentiation-mediated tumour growth and whether it is functionally required for this process is so far unknown.

In this study, we demonstrate that dedifferentiation drives extensive remodelling of the CG niche. This remodelling involves an increase in CG cell numbers, resulting from the combined effects of *de novo* gliogenesis originating from the tumour and the prolonged proliferation of CG cells due to dysregulated ecdysone signalling. The RNA binding proteins Imp and Syncrip (Syp) are temporal factors important for the specification of neuronal diversity as well as tumour malignancy (Narbonne-Reveau *et al*, 2016; Zhu *et al*, 2006). It has been previously shown that the Imp^+^ tumour NBs are the cancer stem cell-like population which determines the persistence and malignancy of the tumour (Narbonne-Reveau *et al*, 2016; Genovese *et al*, 2019). Here, we demonstrate that the less proliferative and more differentiated Syp^+^ tumour NBs are essential for generating ectopic glial cells. Notably, inhibiting Syp disrupts *de novo* gliogenesis, thereby impairing tumour growth. Conversely, the expansion of the glial niche via the expression of Syp or the glial fate determinant Reverse Polarity (Repo) is sufficient to enhance tumour growth. Together, these data show that brain tumours exhibit the remarkable capacity to generate their own niche to optimise conditions for tumour growth. Ways to target the niche hold great promise in restricting brain tumour growth.

## Results

### NB tumours exhibit an expanded cortex glial niche

It is known that the perturbation of asymmetric division caused by the inactivation of the transcription factor Pros in type I NB lineages results in the reversion of GMCs into ectopic NBs (Figure 1 A-B) (Caussinus & Gonzalez, 2005). In this study, we generated a *QUAS-pros^RNAi^* line to inhibit *pros* expression in NB lineages using the QF/QUAS binary gene expression system (Riabinina *et al*, 2015; Potter *et al*, 2010), allowing us to use the GAL4/UAS system to perform gene manipulation in other tissues. Knockdown of *pros* in pan-NB lineages (*elav^C155^-QF2*) resulted in a 21-fold increase in the volume of NBs (see methods), as indicated by the NB marker Dpn (Figure 1 D-F, 120 hours after egg laying, h AEL). It also caused a 1.6-fold increase in proliferative NBs, evidenced by a higher percentage of NBs incorporating 5-ethynyl-2 deoxyuridine (EdU), a nucleoside analogue that is incorporated into newly synthesized DNA of dividing cells, within a 15-minute window (Figure 1 D’–E’ and G). Additionally, tumour induction also resulted in a significant developmental delay (Figure S1 A). Together, our data shows that dedifferentiation induced by *elav-QF2>QUAS-pros^RNAi^* causes brain overgrowth and a delay in pupariation.

To investigate whether dedifferentiation causes changes in the topology of the glial niche, we quantified the number of glial cells using the pan-glial nuclear marker Repo. We observed a significant increase in the total number of glial cells at 120 h AEL in tumour brains compared to controls (Figure 1 D’’-E’’, H). However, glia to NB ratio was reduced compared to the control brain (Figure 1 I), suggesting that while glial expansion occurs in response to tumour growth, it does not keep up with the overall growth of the tumour. A similar expansion of glial cells was observed following the knockdown of *pros* using the pan-NB driver *dnab-GAL4* (Figure S1 B-D). Moreover, the enlargement of the glial cell population seems to be a common characteristic of brain tumors. Type I NB tumours induced by the overexpression of *aPKC* (*aPKC^CAAX^*) (Lee *et al*, 2006) and type II NB tumours resulting from the knockdown of the NHL protein Brat (Betschinger *et al*, 2006) both displayed glial niche expansion (Figure 1 B-C, Figure S1 E-J). This observation suggests that glial niche expansion is a general feature of dedifferentiation-mediated brain tumours.

In the fly brain, glial cells are categorized into surface glia (SG), cortex glia (CG), and neuropil glia (NG) based on their distinct locations within the CNS (Freeman, 2015; Yildirim *et al*, 2019) (Figure 1 J). Surface glia, which include perineural and subperineural glia, are situated at the CNS surface and form the blood-brain barrier (BBB). During larval development, perineural glia proliferate to create a cellular meshwork at the CNS’s outermost layer, directly interacting with the circulating haemolymph. Beneath these, the less proliferative subperineural glia establish the BBB through septate junctions (Stork *et al*, 2008). CG enwraps NBs and their neuronal lineages serving as their direct niche. These glial cells undergo extensive remodelling during larval development, including processes such as membrane extension, cell division, endoreplication, and cell fusion (Pereanu *et al*, 2005; Avet-Rochex *et al*, 2012; Rujano *et al*, 2022; Spéder & Brand, 2018). We discovered that the increase in glial cell numbers associated with *pros^RNAi^*-mediated tumour growth is primarily attributed to an expansion of CG cells (Richier *et al*, 2017), marked by the expression of *wrapper-GAL4* (Figure 1 K-M); with little or no increase in perineural glial cells marked by *GMR85G01-GAL4* (Kremer *et al*, 2017) (Figure 1 N-P). Consistent with previous findings (Banach-Latapy *et al*, 2023), we observed that the induction of the *pros^RNAi^* tumour, resulted in CG chambers containing multiple NBs instead of enwrapping individual NBs and their neuronal progeny (Figure S1 K-L’). Furthermore, these CG cells were significantly smaller in the tumour brains compared to those in the control brains (Figure S1 M-O).

### CG cells proliferate for longer but not faster in tumour brains

Next, we assessed how the extra glial cells arose in the tumour brains. We first tested if glial cells proliferated faster in tumour brains compared to the control. After 22 h of EdU feeding, we detected a 1.3-fold increase in the number of EdU^+^ glial cells compared to control, suggesting that cumulatively, more glial cells were generated in the tumour brains (Figure 2 A-D). However, the percentage of glial cells that were EdU^+^ was not significantly altered in the tumours versus the control (Figure 2 E), suggesting that the rate of glial proliferation was not significantly different between control and tumour brains.

**Figure 2.**
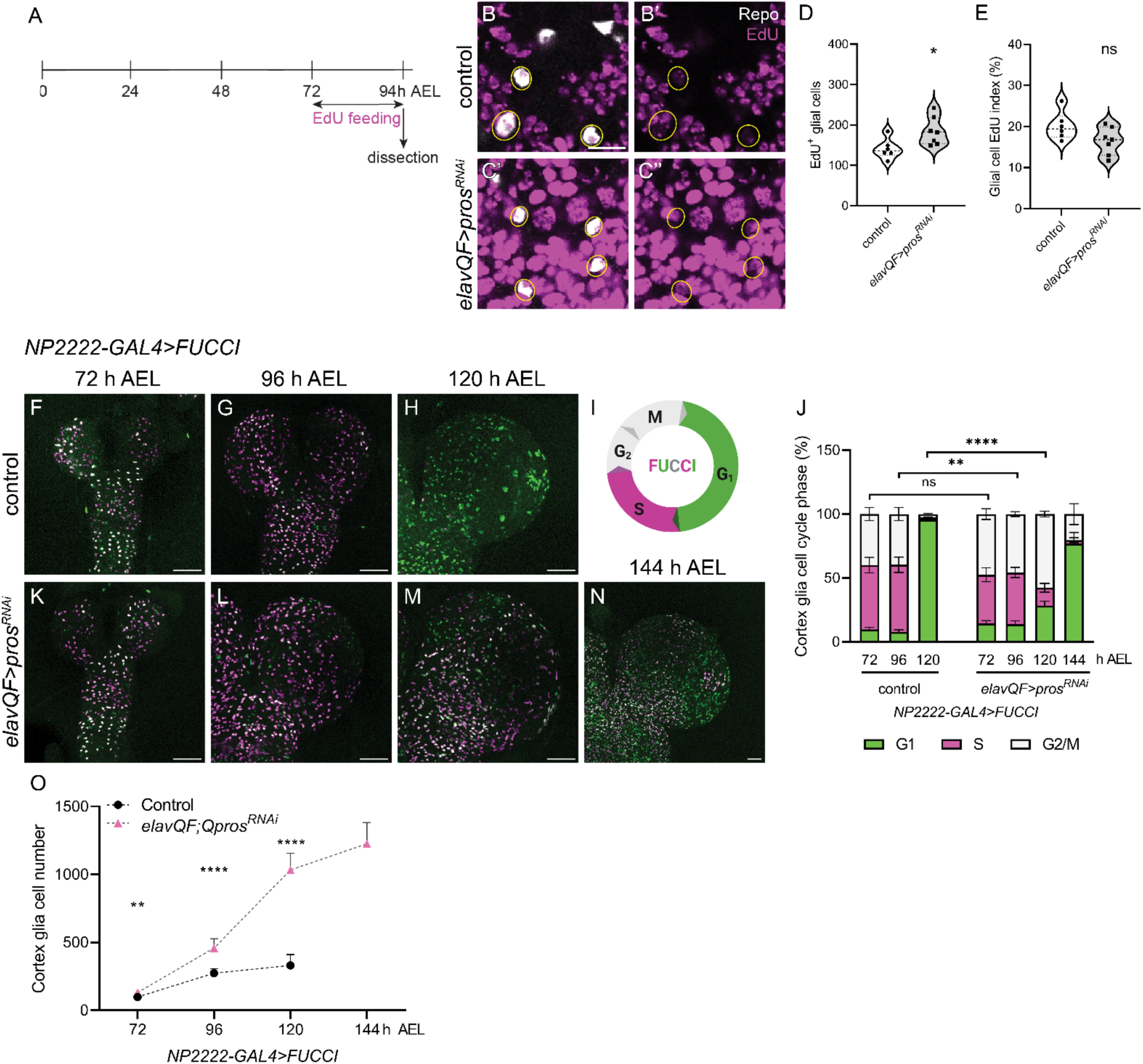
Cortex glial (CG) cells proliferate for longer, but not faster, in tumour brains. A. Schematic depicting the experiment setup: larvae were fed with food supplemented with EdU from 72 h AEL, and brains were dissected at 94 h AEL. B-C’’. Representative images showing that there are more EdU^+^ cells (Repo^+^) (yellow circles) in *elavQF>pros^RNAi^* brains compared to control brains. Scale bar= 10 μm. D. Quantification of the number of EdU^+^ glial cells (Repo^+^) in B and C, performed using an unpaired t-test. Control: n= 6, m= 140.7 ± 10.16. *pros^RNAi^*: n= 7, m= 184.4 ± 13.19. *p < 0.05. E. Quantification of the % glial EdU index (Repo^+^ EdU^+^ cells/ Repo^+^ cells) in B and C, performed using an unpaired t-test. Control: n= 6, m= 20.12 ± 1.4. *pros^RNAi^*: n= 7, m= 16.44 ± 1.25. ns (not significant) p > 0.05. F-H. Representative images (maximum projections) showing the cell cycle stages of CG cells visualised by *NP2222-GAL4>FUCCI* in control brains at 72, 96 and 120 h AEL. Scale bar= 50 μm. I. Schematic depicting FUCCI marking cells in G1 phase in green, cells in S phase in magenta, and cells in G2/M phase in white. K-N. Representative images (maximum projections) showing the cell cycle stages of CG cells visualised by *NP2222-GAL4>FUCCI* in *elavQF>pros^RNAi^* tumour brains at 72, 96, 120 and 144 h AEL. Scale bar= 50 μm. J. Quantification of the cell cycle phase distribution of CG in F-N, performed by Chi-square test. See Table 3 for n, mean, SEM, and adjusted p-values. ns (not significant) p > 0.05, **p < 0.01, ****p < 0.0001. O. Quantification of the number of CG cells at 72, 96, 120 and 144 h AEL in control and *elavQF>pros^RNAi^* tumour brains, as marked by cells expressing *NP2222-GAL4>FUCCI* in F-N. Quantifications performed using a Mann-Whitney test (72 h AEL) and unpaired t-test (96 and 120 h AEL). See Table 4 for n, mean, SEM, and adjusted p-values. **p < 0.01, ****p < 0.0001.

To gain a deeper understanding of the cell cycle state of CG cells, we employed the Fly-FUCCI system (Figure 2 I) (Zielke *et al*, 2014), driven by the CG-specific driver *NP2222-GAL4*. FUCCI marks cells in the G1 phase with ECFP (green, Figure 2 F-N), in S phase with Venus (magenta, Figure 2 F-N), and in G2 and M phases with ECFP and Venus (white, Figure 2 F-N). In control brains, most of the CG cells are actively cycling (in S or G/M phase) until 120 h AEL (Figure 2 F-H) (Banach-Latapy *et al*, 2023), whereas CG cells in tumour brains are still dividing at 144 h AEL (Figure 2 K-N, J). Although the overall period of CG proliferation was significantly prolonged in tumour brains, the rate of CG proliferation was either not significantly different at 72 h AEL or slightly slower at 96 h AEL, as indicated by the distribution of CG cells in G1, S, and G2/M phases (Figure 2 J). Interestingly, although CG cells in tumour brains did not proliferate faster, we observed an increase in the total number of CG cells in tumour brains compared to controls from 72 h AEL (Figure 2 O).

### Ecdysone signalling regulates the termination of CG proliferation

It is well known that at the end of larval stages, a surge in the levels of the steroid hormone ecdysone trigger pupa formation (Riddiford, 1993). Tumour-bearing larvae exhibited delayed pupariation (Figure S1 A) suggesting a potential reduction in the systemic levels of ecdysone. To investigate whether ecdysone levels were diminished in tumour-bearing animals, we conducted quantitative RT-PCR to measure the expression levels of *phantom* (*phm*) and *spookier* (*spok*), key genes involved in ecdysone biosynthesis. Compared to control animals, tumour-bearing larvae showed a reduction in the transcription of *spok* and a decrease (though not statistically significant) in the expression levels of *phm* (Figure 3 A), which may account for the developmental delay observed (Figure S1A).

**Figure 3.**
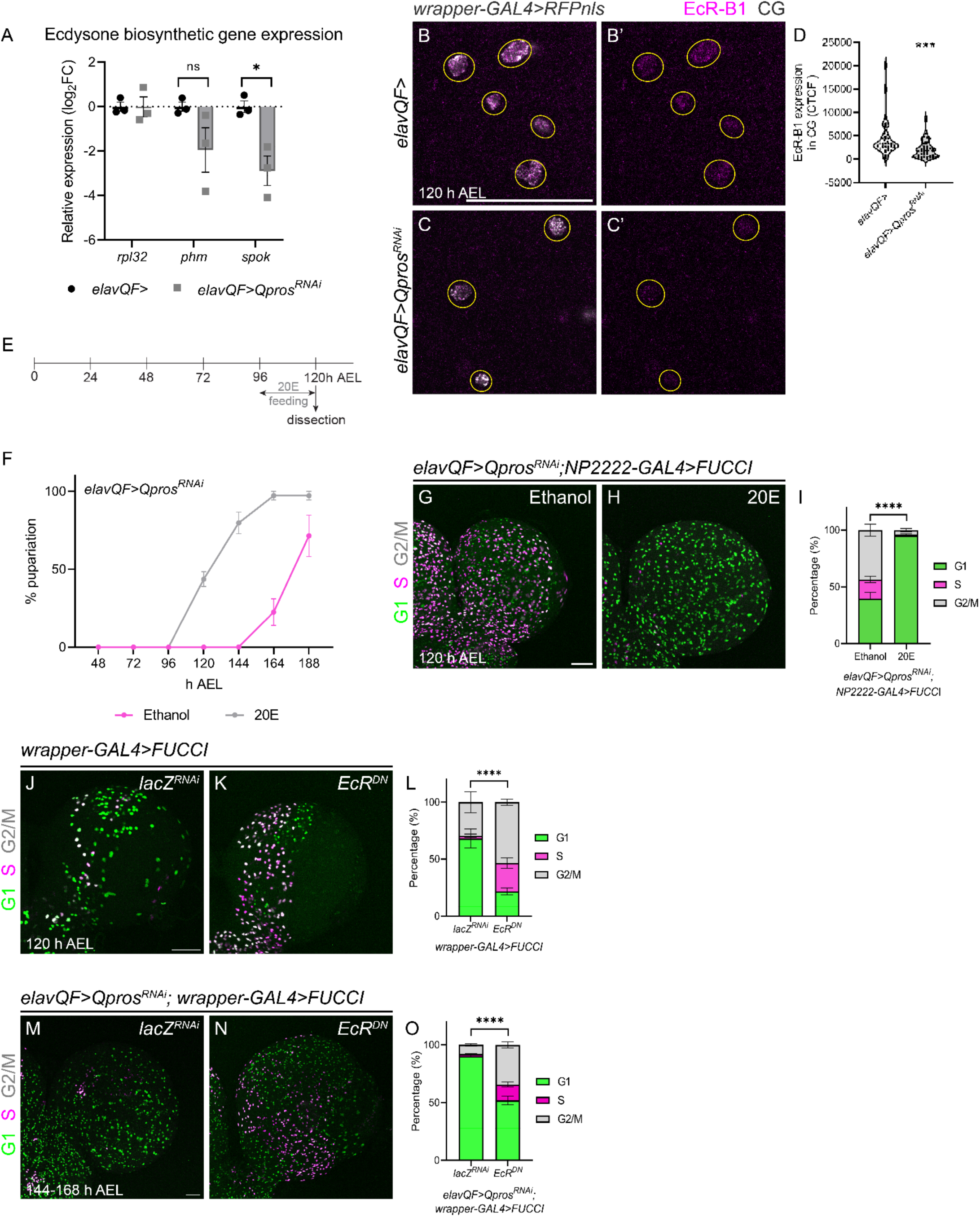
Ecdysone signalling controls the cessation of cortex glia (CG) proliferation. A. Expression levels of ecdysone biosynthetic genes *phantom* (*phm*) and *spookier* (*spok*) in control (*elavQF>+*) and tumour-bearing animals (*elavQF>pros^RNAi^*) at 120 h AEL, relative to *rpl32* expression. Quantifications performed by independent student t-tests. *phm - elavQF*: n= 3, m= 0 ± 0.2. *elavQF*>*pros^RNAi^*: n= 3, m= −1.95 ± 1. *spok - elavQF*: n= 3, m= 0 ± 0.26. *elavQF*>*pros^RNAi^*: n= 3, m= −2.88 ± 0.66. *p < 0.05, ns (not significant) p > 0.05. B-C’. Representative images showing EcR-B1 staining in control (*elavQF>+*) and *elavQF>pros^RNAi^* brains at 120 h AEL in CG cells marked by *wrapper-GAL4>RFP.nls* (circled). EcR-B1 is expressed at higher levels in CG of control brains compared to *elavQF>pros^RNAi^*. Scale bar= 50 μm. D. Quantification of EcR-B1 expression levels in B-C’, performed using a Mann-Whitney test. *elavQF*: n= 40, m= 4259 ± 586.7. *elavQF>Qpros^RNAi^*: n= 40, m= 2075 ± 283.2. ***p < 0.001. E. Schematic depicting the experimental set-up in F and G-I. Larvae were fed with standard fly food supplemented with ethanol or 20E in ethanol for 24 h from 96 h AEL and dissected at 120 h AEL. F. Feeding *elavQF>Qpros^RNAi^* larvae with 20E results in their earlier pupariation, in comparison to controls fed with ethanol. Statistical analysis performed using a two-way ANOVA followed by Šídák’s multiple comparisons post-hoc test. See Table 5 for n, mean, SEM, and adjusted p-values. G-H. Representative images showing the cell cycle stages of CG cells visualised by *NP2222-GAL4>FUCCI* in tumour-bearing animals fed with ethanol vs. 20E, at 120 h AEL. Scale bar= 50 μm. I. Quantification of the cell cycle phase distribution of CG in G-H. Statistical analyses using Chi-square test (****p < 0.0001) and two-way ANOVA followed by Šídák’s multiple comparisons post-hoc test. See Table 6 for n, mean, SEM, and adjusted p-values. J-K. Representative images showing the cell cycle stages of CG cells visualised by *wrapper-GAL4>FUCCI* where *lacZ^RNAi^* or *EcR^DN^* was specifically expressed in CG, at 120 h AEL. Scale bar= 50 μm. L. Quantification of the cell cycle phase distribution of CG in J-K. Statistical analyses using Chi-square test (****p < 0.0001) and two-way ANOVA followed by Šídák’s multiple comparisons post-hoc test. See Table 7 for n, mean, SEM, and adjusted p-values. M-N. Representative images (maximum projections) showing the cell cycle stages of CG cells visualised by *wrapper-GAL4>FUCCI* in *elavQF>pros^RNAi^* tumour brains where *lacZ^RNAi^* or *EcR^DN^* was specifically expressed in CG, at 144-168 h AEL. Scale bar= 50 μm. O. Quantification of the cell cycle phase distribution of CG in M-N, Statistical analyses using Chi-square test (****p < 0.0001) and two-way ANOVA followed by Šídák’s multiple comparisons post-hoc test. See Table 8 for n, mean, SEM, and adjusted p-values.

As high levels of ecdysone are required to promote the cessation of neurogenesis, it is therefore possible that the reduced ecdysone signalling in CG of tumour brains could account for their prolonged persistence in the cell cycle. Ecdysone signalling is transduced via the binding of ecdysone to the ecdysone receptors (EcRs, including the isoforms EcR-A1 and B1) and its co-receptor Ultraspiracle (USP) (Colombani *et al*, 2005). While EcR-B1 was detected in the nuclei of control CG cells at 120 h AEL, its expression level was significantly downregulated in CG cells of tumour brains (Figure 3 B-D). Therefore, low EcR-B1 expression in CG of tumour brains could account for reduced ecdysone signalling, and in turn delay CG cell cycle exit.

Next, we asked if increasing global ecdysone levels could trigger precocious cessation of CG proliferation. We fed tumour-bearing animals with the active form of ecdysone, 20 hydroxy-ecdysone (20E) from 96 h AEL (Figure 3 E), and found this manipulation was able to cause timely pupariation of tumour-bearing animals (Figure 3 F). Consistent with previous reports in the wing, where high levels of 20E triggered proliferation arrest (Strassburger *et al*, 2021; Parker & Struhl, 2020; Perez-Mockus *et al*, 2023), we found that CG cells in tumour-bearing animals fed with 20E ceased divisions at 120 h AEL (most of the glial cells are in G1, Figure 3 G-I), whereas CG cells from ethanol-fed control animals continued to divide at this time point (Figure 3 G-I). Together, this data shows that ecdysone signalling is sufficient to terminate CG proliferation.

Using the CG-specific driver *wrapper-GAL4*, we analysed the cell cycle profile of CG cells in wild-type (WT) brains. At 120 h AEL, we found that most CG cells were in the G1 phase, approximately 25% were in G2/M, and only a small fraction were in the S phase of the cell cycle, indicating they were undergoing cell cycle exit (Figure 3 J and L). To determine whether CG cells in WT brains stop dividing at this time point due to their response to ecdysone through EcR, we overexpressed a dominant-negative form of EcR (*EcR^DN^*) (Cherbas *et al*, 2003) using *wrapper-GAL4*, to block ecdysone signalling. We observed that this intervention significantly increased the proportion of CG cells in the S and G2/M phases at 120 h AEL (Figure 3 J-L), suggesting that blocking ecdysone signalling allows CG cells to continue proliferating. Supporting the hypothesis that ecdysone signalling suppresses CG proliferation in WT animals, blocking ecdysone signalling in CG cells of tumour-bearing animals through *EcR^DN^* expression also effectively extended the overall duration of CG proliferation. This was indicated by a significant increase in the percentage of CG cells in S and G2/M phases of the cell cycle (Figure 3 M-O) at 144-168 h AEL (tumour animals remain as larvae due to developmental delay). Collectively, our findings indicate that ecdysone signalling plays a role in regulating CG termination and that reduced ecdysone levels in tumour-bearing animals leads to prolonged proliferation of CG cells.

### WT and Pros loss-of-function type I NBs generate ectopic glial cells

While the extended period of CG proliferation in tumour brains may contribute to the overall increase in CG cell number, it cannot account for the substantial rise in CG cell number observed at 72, 96, and 120 h AEL in tumour brains (Figure 2 O). We therefore hypothesised that the excess CG cells were generated by other progenitor-like populations. It is known that a subset of embryonic glial cells are generated by neuroglioblasts, common precursors of both neurons and glial cells (Pereanu *et al*, 2005). Furthermore, in the larval CNS, it has been reported that both type II NB lineages as well as medulla NBs in the OLs are capable of generating glial cells in addition to neurons (Syed *et al*, 2017; Viktorin *et al*, 2011, 2013; Li *et al*, 2013). Interestingly, upon the reduction of *pros^RNAi^* tumour growth, via the inhibition of the FGF (Heartless, Htl) (Beiman *et al*, 1996), EGF (dSor) (La Marca *et al*, 2023) or PI3K (UAS-Pten) (Goberdhan *et al*, 1999) signalling pathways, glial cell numbers were significantly reduced (Figure S2 A-E), suggesting it is possible that the growth of NB-derived tumours is coupled with the production of ectopic glia.

To determine whether these glial cells originate from type I NB lineages and identify the cell of origin within *pros^RNA^*^i^-induced tumours, we conducted lineage tracing studies using G-TRACE (Figure 4B). This technique reports both current and historical GAL4 expression (Evans *et al*, 2009), allowing us to track cell fate changes within *dnab-GAL4*-labeled NB lineages. *dnab-GAL4>UAS-GFP* marks NB lineages (marked by a dotted yellow line, Figure S2 F-F’’’) encompassing NBs (Dpn^+^), neurons, but not glial cells (Repo^+^) (Figure S2 F-F’’’). Similarly, through G-TRACE analysis, we found that Repo^+^ glial cells did not coincide with the real-time *dnab-GAL4* expression (magenta) in either control or *pros^RNAi^*-induced tumours (Figure 4 C-D, dotted yellow circles). However, the Repo*^+^* glial cells were found to overlap with the historical expression of *dnab-GAL4* (green) in both control and tumor brains (Figure 4 C-D’’’’, yellow circles), indicating that these glial cells likely originated from NB lineages. Additionally, a higher number of Repo*^+^* cells with historical *dnab-GAL4* expression was observed in the tumour compared to the control (Figure 4 E). This suggests that tumour-derived glial cells may contribute to the overall increase in glial cell numbers within tumour brains. These tumour-derived glial cells were located throughout the brain, both on the ventral side of the brain lobes (Figure 4 A and C-D’’’’) as well as in the VNC, where only type I NBs can be found (Figure S2 G-G’’’’).

**Figure 4.**
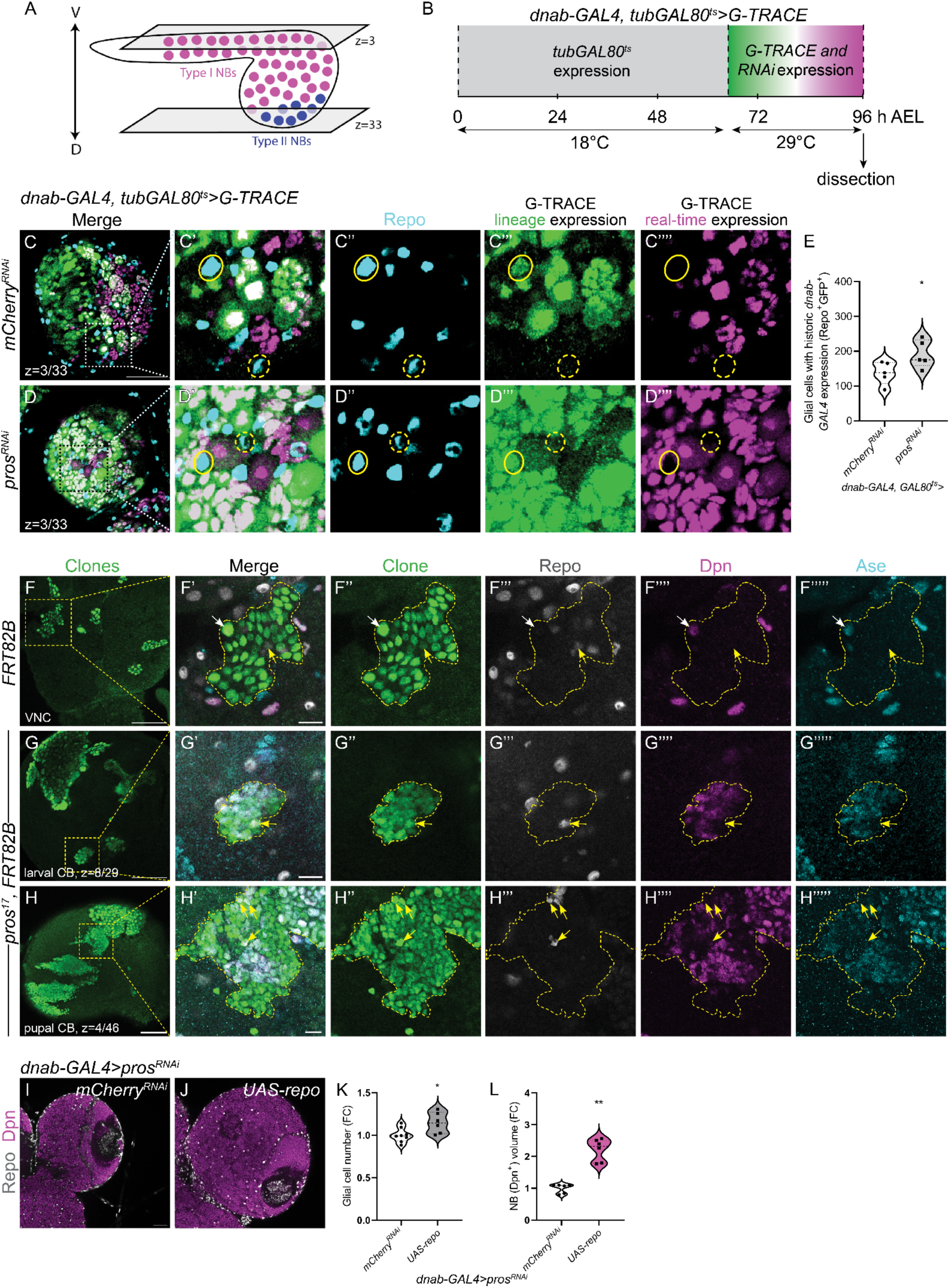
Tumour neuroblasts (NBs) generate glial cells. A. Schematic depicting the ventral (V)-dorsal (D) axis of the *Drosophila* larval brain. Rectangular boxes represent the plane of vision throughout the stack. Z-stack = 3 is in the V side of the brain, where only type I NBs are found. Z-stack = 33 is in the dorsal side of the brain, where type II NBs are found. B. Schematic depicting the experimental set-up in C-D’’’’. Larvae were kept at 18 °C until 72 h AEL and then moved to 29 °C, allowing G-TRACE and RNAi expression for 24 h before dissection at 96 h AEL. C-D’’’. Representative images showing NB lineages (*dnab-GAL4*) expressing G-TRACE in control (*mCherry^RNAi^*) or tumour brains (*pros^RNAi^*). Real-time expression is visualised in magenta, lineage expression in green and glial cells in cyan. C’-C’’’’ and D’-D’’’’ are zoomed-in images from C and D, respectively. Yellow circles mark glial cells with G-TRACE lineage expression and yellow dashed circles mark glial cells with no G-TRACE lineage expression. Scale bar= 50 μm. E. Quantification of the number of glial cells (Repo^+^) in C and D with historic *dnab-GAL4* expression, performed using an unpaired t-test. *mCherry^RNAi^*: n= 5, m= 137.4 ± 14.55. *pros^RNAi^*: n= 5, m= 191.2 ± 17.77. *p < 0.05. F-H. Representative images showing glial cells (Repo^+^) within WT (*FRT82B*) and *pros^17^* MARCM clones (GFP), which do not express NB markers Dpn and Ase. Dotted lines outline the clone, white arrows point to NBs and yellow arrows point to glial cells within the clone. F-F’’’’’: WT clone within the larval VNC. G-G’’’’’: *pros^17^* clone in the ventral side of the larval CB. H-H’’’’’: *pros^17^* clone in the ventral side of the pupal CB. Scale bar= 50 μm (in F, G, H), 10 μm (in F’, G’, H’). I-J. Representative images of brain lobes showing that *UAS-repo* in *dnab-GAL4>pros^RNAi^*lineages caused increased glial cell (Repo) number and NB volume (Dpn) compared to the *mCherry^RNAi^* control. Scale bar= 50 μm. K. Quantification of glial cell number (Repo^+^, FC) in I-J, performed using an unpaired t-test. Data normalised to controls. *mCherry^RNAi^*: n= 8, m= 1 ± 0.03. *UAS-repo*: n= 6, m= 1.145 ± 0.05. *p < 0.05. L. Quantification of NB volume (Dpn^+^, FC) in I-J, performed using a Mann-Whitney test. Data normalised to controls. *mCherry^RNAi^*: n= 6, m= 1 ± 0.06. *UAS-repo*: n= 6, m= 2.215 ± 0.14. **p < 0.01.

Next, we induced MARCM clones, which also allowed us to investigate the identity of the progeny cells derived from NBs. WT clones in the VNC are originated from type I NBs (Dpn^+^ Ase*^+^*, white arrow, Figure 4 F-F’’’’’) (Homem & Knoblich, 2012). Consistent with our lineage tracing results, we found that these WT clones produced sporadic Repo*^+^*glial cells at 120 AEL, as previously shown by (Pereanu *et al*, 2005) (yellow arrow, Figure 4 F-F’’’’’). Repo^+^ cells were also identified in *pros* mutant clones (*pros^17^*) on the ventral surface of larval and pupal CBs where type I NBs are located (yellow arrows, Figure 4 G-H’’’’’) (Homem & Knoblich, 2012). Notably, in both WT and *pros^17^* clones, Repo^+^ glial cells did not express the NB markers Dpn or Ase (Figure 4 F-H, yellow arrows), indicating that these cells had differentiated into glial cells. Together, our clonal and lineage studies suggest that both type I NBs and Pros loss-of-function type I NBs are capable of generating glial cells.

Next, we investigated the factors that might promote gliogenesis in these lineages. A subset of embryonic NBs, known as neuroglioblasts, has the ability to generate both neuronal and glial progeny (Bossing *et al*, 1996; Schmidt *et al*, 1997; Schmid *et al*, 1999; Akiyama *et al*, 1996). Gliogenesis occurs through transient and low expression of the glial master regulator glial cells missing (*gcm*) (Hosoya *et al*, 1995; Jones *et al*, 1995; Vincent *et al*, 1996; Freeman & Doe, 2001). Gcm is a transcriptional regulator directly controlling *repo*, which promotes glial differentiation (Campbell *et al*, 1994; Halter *et al*, 1995; Xiong *et al*, 1994; Akiyama *et al*, 1996). We next assessed if these transcription factors were necessary for promoting glial fate in WT and *pros^17^*clones. We first used *gcm-lacZ* to visualize the expression pattern of *gcm*. While we did not observe any *gcm-lacZ* expression in WT clones (Figure S2 H-H’’’’), we detected some expression in *pros^17^*mutant clones (Figure S2 I-I’’’’, yellow arrows). Interestingly, *gcm-lacZ* expression never colocalized with the NB marker Dpn or the glial marker Repo (Figure S2 I-I’’’’). We hypothesized that this could be because *gcm* is expressed only transiently (Freeman & Doe, 2001). Next, we tested the necessity of *gcm* to promote gliogenesis in *pros^RNAi^* tumours. The expression of a dominant negative form of *gcm* (*gcm^DN^*) in tumour lineages did not significantly alter glial cell number or tumour size (Figure S3 A-D), suggesting that Gcm is not required for tumour-derived gliogenesis. Given the transient nature of *gcm* expression, we next investigated the role of *repo* in promoting gliogenesis in *pros^RNAi^* tumours. Upon the expression of an RNAi against the mature glial transcription factor *repo* in tumour lineages from 48 h AEL, we observed a slight decrease in tumour size but no significant decrease in glial cell number compared to controls (Figure S3 E-H). In contrast, overexpression of *repo* within tumour lineages led to an increase in both glial cell number and tumour volume (Figure 4 I-L). These findings suggest that while *gcm* and *repo* are not required for tumour gliogenesis, *repo* is sufficient to drive glial cell expansion. We next investigated whether *repo* was sufficient to promote gliogenesis and increase the number of NBs in the WT brain. However, in contrast to the tumour brains, *repo* overexpression in *dnab-GAL4* lineages did not result in an increase in either glial cell number or type I NB number (Figure S3 I-L), indicating that the role of *repo* in promoting gliogenesis is specific to tumour NBs.

Altogether, our findings reveal a role for *repo* in promoting gliogenesis in tumour NBs and demonstrate that expanding the glial niche by driving glial cell fate and increasing glial cell numbers is sufficient to enhance tumour growth. This suggests that the expanded niche may provide support for the growth of NB tumours.

### Ectopic glial cells are produced in the Syp^+^ temporal window

In all NBs, temporal patterning is regulated by cascades of temporal transcription factors (tTFs). tTFs are deployed to create cellular diversity and determine the commencement and the cessation of NB proliferation (Bayraktar & Doe, 2013). RNA-binding proteins Imp and Lin-28 are detected in young type I NBs prior to 84 h AEL (Liu *et al*, 2015), whereas the RNA-binding protein Syncrip (Syp) is expressed in older type I NBs from ∼88 h AEL till their termination at around 24 h after pupal formation (Syed *et al*, 2017). Neurons generated during the early temporal window (Imp^+^) express the TF Chinmo, whereas those generated during the late temporal window (Syp^+^) express the TF Broad (Br) (Liu *et al*, 2015; Maurange *et al*, 2008). Furthermore, late temporal gene expression is essential for neuronal and glial cell type specification in type II NBs (Syed *et al*, 2017). Previously, it was shown that *pros* mutant clones induced during the early Imp^+^ temporal window exhibit malignant growth potential (Narbonne-Reveau *et al*, 2016), whereas clones induced during the late Syp^+^ window disappear upon the differentiation of tumour cells (Genovese *et al*, 2019). To assess during which time window the type I tumour NBs give rise to glial cells, we performed *pros* knockdown (*dnab-GAL4*) for 48 h, either during the early Imp^+^Chinmo^+^ time window (24-72 h AEL) or during the late Syp^+^Br^+^ time window (72-120 h AEL) (Figure 5 A). We observed an increase in glial cell numbers when tumours were induced in the late temporal window, but not during the early temporal window (Figure 5 B-F). To investigate the function of Syp in tumour gliogenesis, we knocked it down in *dnab-GAL4>pros^RNAi^* tumours from 48 h AEL. *syp* knockdown led to a reduction in glial cell number (Figure 5 G-I). Interestingly, this manipulation also caused a significant reduction in tumour size (Figure 5 G-H, J). This contrasted with a previous report, where it was shown that *syp* knockdown in *Poxn>pros^RNAi^* tumours (*Poxn-GAL4* is specifically expressed in six thoracic NB clusters) caused an increase in tumour size by adult stages (Genovese *et al*, 2019). In our hands, the knockdown of *syp* (using two independent RNAi lines) in *Poxn>pros^RNAi^* tumours did not cause significant changes to tumour size by 120 h AEL (Figure S4 A-D). To understand the differential effects of *syp* knockdown in *dnab* vs. *Poxn* lineages, we examined the cellular composition of the *Poxn* lineages. We found that, in contrast to other type I lineages, *Poxn* lineages do not produce ectopic glial cells (Figure S4 E-F). Therefore, it is possible that Syp only exert its effect on lineages that produce glial cells. Next, we tested if extending the temporal window for glial cell production in *dnab-GAL4*-induced tumour lineages would significantly increase glial cell numbers. *syp* overexpression in *dnab>pros^RNAi^* tumours led to a significant increase in both glial cell number and tumour size (Figure 5 K-N). This suggests that similar to the previous experiment in Figure 4 I-L, where *repo* overexpression in the tumour promotes gliogenesis and tumour growth, *syp* overexpression promotes gliogenesis from the tumour, contributing to the formation of a glial niche that supports tumour growth. Next, we investigated whether *syp* expression is required and sufficient to drive gliogenesis in WT type I NBs. Knockdown of *syp* using *dnab-GAL4* from 48 h AEL did not affect either NB or glial cell numbers (Figure S4 G-J). Similarly, overexpression of *syp* in *dnab-GAL4* lineages also did not alter NB or glial cell numbers (Figure S4 K-N), suggesting that the necessity of Syp to generate ectopic glial cells is specific to tumour NBs. Together, our data suggest that glial cells are produced during the *Syp^+^* temporal window in type I tumour lineages, except in certain lineages, such as *Poxn* lineages, which do not generate ectopic glial cells. Moreover, Syp is necessary and sufficient for the *de novo* gliogenesis, which results in the expansion of the glial niche and in turn sustains tumour growth.

**Figure 5.**
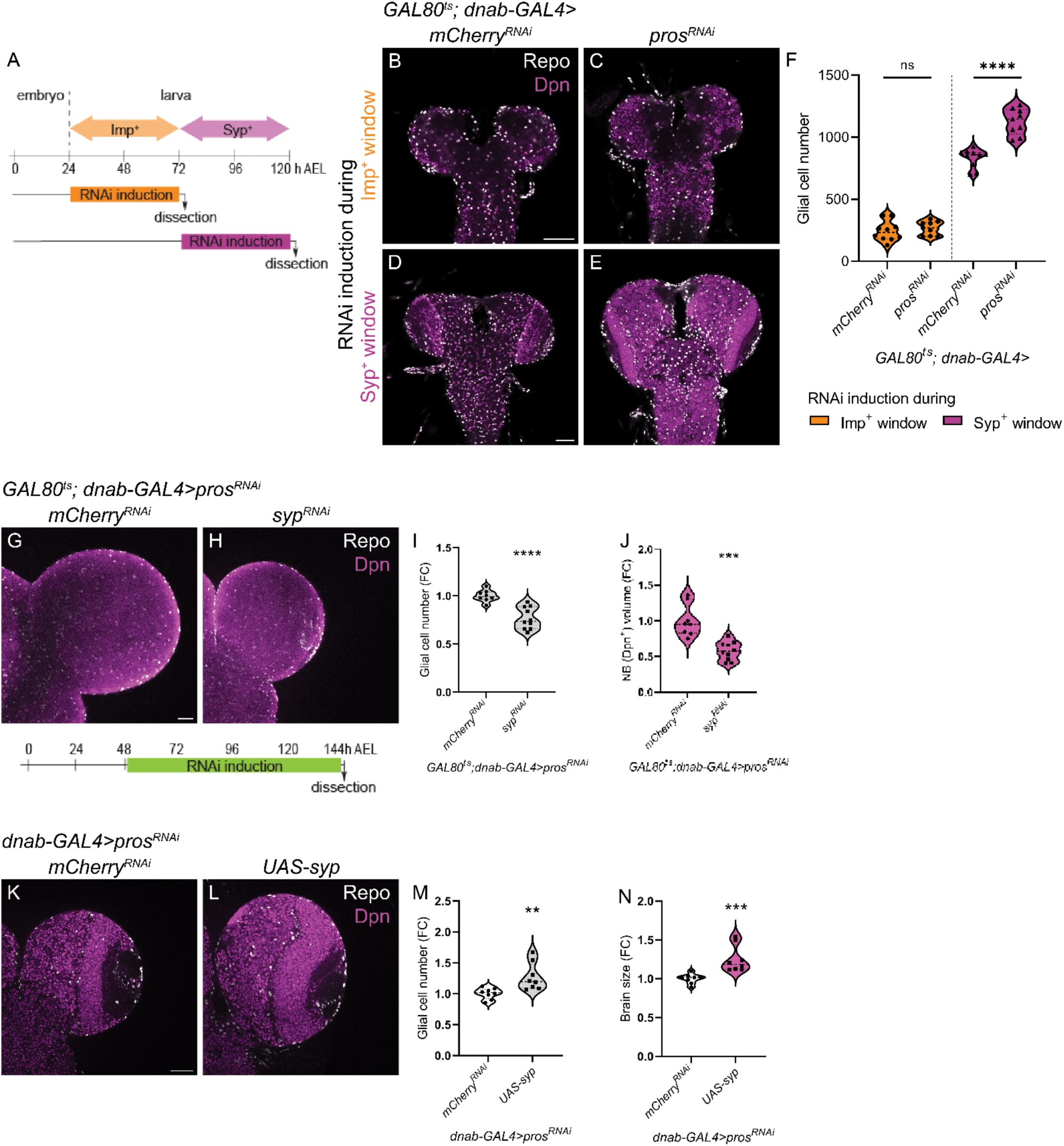
Ectopic glial cells are produced in the Syp^+^ temporal window. A. Schematic depicting the experimental set-up in B-E. *pros* RNAi induction using *GAL80^ts^;dnab-GAL4* was carried out either during the Imp^+^ window from 24-72 h AEL, or during the Syp^+^ window from 72-120 h AEL. B-E. Representative images from brain lobes where *pros^RNAi^* was induced during either the Imp^+^ or the Syp^+^ temporal window. NBs were marked by Dpn, and glial cells by Repo. Scale bar= 50 μm. F. Quantification of glial cell number (Repo^+^) from B-E. Unpaired t-test performed for RNAi induction during Imp^+^ window - *mCherry^RNAi^*: n= 10, m= 243.1 ± 25.22. *pros^RNAi^*: n= 9, m= 266.7 ± 18.87. Mann-Whitney test performed for RNAi induction during Syp^+^ window - *mCherry^RNAi^*: n= 10, m= 832.1 ± 23.91. *pros^RNAi^*: n= 9, m= 1126 ± 34.89. ns (not significant) p > 0.05, ****p < 0.0001. G-H. Representative images of brain lobes, where *mCherry^RNAi^* or *syp^RNAi^* were induced from 48 h AEL using *GAL80^ts^;dnab-GAL4*>*pros^RNAi^* and dissected at 144 h AEL. NBs were marked by Dpn, and glial cells by Repo. Scale bar= 50 μm. I. Quantification of glial cell number (Repo^+^, FC) in G-H, performed using an unpaired t-test. Data normalised to controls. *mCherry^RNAi^*: n= 8, m= 1 ± 0.02. *syp^RNAi^*: n= 10, m= 0.77 ± 0.4. ***p < 0.001, ****p < 0.0001. J. Quantification of NB volume (Dpn^+^, FC) in G-H, performed using an unpaired t-test. Data normalised to controls. *mCherry^RNAi^*: n= 8, m= 1 ± 0.02. *syp^RNAi^*: n= 10, m= 0.77 ± 0.4. ***p < 0.001, ****p < 0.0001. K-L. Representative images of brain lobes, where *mCherry^RNAi^* or *UAS-syp* were induced from 0 h AEL in *dnab-GAL4>pros^RNAi^* tumour lineages. Scale bar= 50 μm. M. Quantification of glial cell number (Repo+, FC) in K-L, performed using an ordinary one-way ANOVA. The *mCherry^RNAi^* control is the same as in Figure S3 A and D. *mCherry^RNAi^*: n= 8, m= 1 ± 0.03. *UAS-syp*: n= 8, m= 1.27 ± 0.08. **p < 0.01. N. Quantification of brain size (FC) in K-L, performed using a Kruskal-Wallis test. The *mCherry^RNAi^* control is the same as in Figure S3 A and C. *mCherry^RNAi^*: n= 8, m= 1 ± 0.02. *UAS-syp*: n= 8, m= 1.25 ± 0.06. ***p < 0.001.

### The CG niche sustains NB tumour growth

Having demonstrated that promoting glial niche expansion (via *UAS-repo* or *UAS-syp*) can enhance tumour growth, and that inhibition of gliogenesis (via *syp^RNAi^*) can perturb tumour growth, we next explored if disrupting the niche via other means could also affect tumour growth. As both proliferation and endoreplication have been implicated in the expansion of the CG niche (Rujano *et al*, 2022), we investigated if these processes are required in CG to accommodate the growth of the tumour. We inhibited glial cell cycle progression through the expression of *dacapo* (*dap*) (Lane *et al*, 1996) and *retinoblastoma-family protein* (*Rbf*) (Xin *et al*, 2002) using the pan-glial driver *repo-GAL4* with a temperature-sensitive *GAL80^ts^*, where the expression of the transgenes was induced at 48 h AEL to bypass embryonic development. This manipulation was sufficient to reduce glial volume and tumour size (Figure 6 A-D). This reduction occurred via reduced NB proliferation as indicated by reduced EdU incorporation (Figure 6 E), without significantly affecting the differentiation status of the tumour (percentage of Elav^+^ neuron volume/DAPI^+^ brain volume, Figure 6 F). Similarly, inhibition of *double parked* (*dup*), known to regulate glial cell endoreplication (Rujano *et al*, 2021), in CG cells with *nrv2-GAL4,* significantly reduced both glial number as well as tumour size (Figure 6 G-J). This result was striking because, in the WT brain, although *dup* knockdown reduced glial cell number (Rujano *et al*, 2021), it did not affect NB number (Figure S5 A-D). This suggests that tumour NBs are more dependent on their glial niche than their WT counterparts. Furthermore, the expression of a RNAi against *m*yc, a global growth regulator (Johnston *et al*, 1999), also reduced both parameters (Figure S5 E-H). Finally, when we induced cell death in cortex glial cells by temporally activating the apoptotic genes *hid* and *rpr* during the late stages of larval development, from 96 h AEL to 144 h AEL, we observed a significant reduction in both glial cell number and tumour growth (Figure 6 K-N). Altogether, these data suggest that the glial niche is essential for sustaining tumour growth.

**Figure 6.**
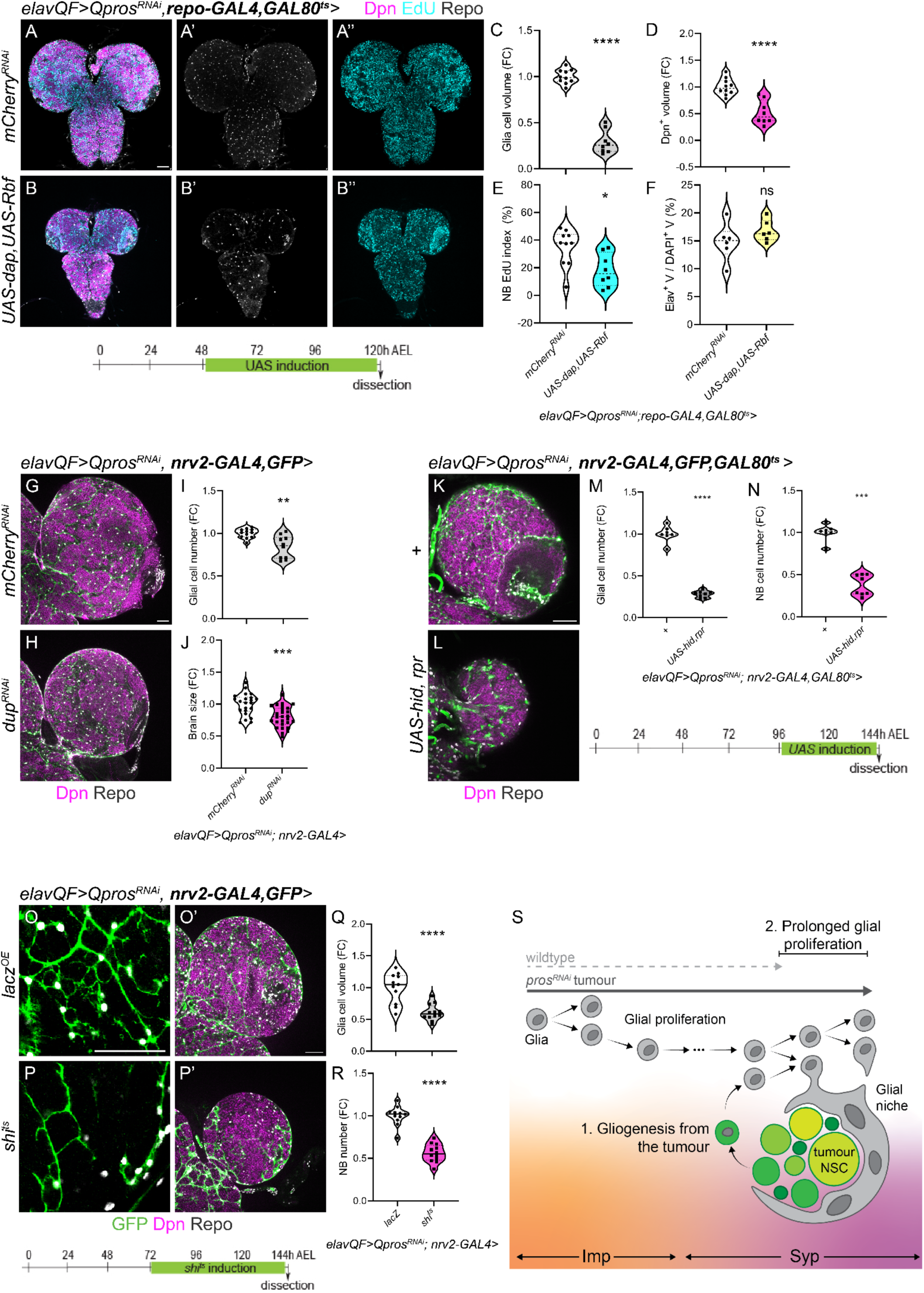
Disruptions of the cell cycle, endoreplication or dynamin activity in the CG inhibit tumour growth. A-B’’. Representative images of brain lobes, where *mCherry^RNAi^* or *UAS-Dap,UAS-Rbf* were induced in glial cells of tumour-bearing animals from 48 h AEL using *elavQF>Qpros^RNAi^*, *repo-GAL4,GAL80^ts^* and dissected at 120 h AEL. NBs were marked by Dpn, and glial cells by Repo. *in vitro* EdU incorporation was performed after dissection for 15 mins. Scale bar= 50 μm. C. Quantification of glial cell volume (Repo^+^, FC) in A-B’’, performed using an unpaired t-test. Data normalised to controls. *mCherry^RNAi^*: n= 10, m= 1 ± 0.02. *UAS-Dap,UAS-Rbf*: n= 8, m= 0.29 ± 0.04. ****p < 0.0001. D. Quantification of NB volume (Dpn^+^, FC) in A-B’’, performed using an unpaired t-test. Data normalised to controls. *mCherry^RNAi^*: n= 10, m= 1 ± 0.05. *UAS-Dap,UAS-Rbf*: n= 8, m= 0.52 ± 0.08. ****p < 0.0001. E. Quantification of NB EdU index (%) in A-B’’, performed using an unpaired t-test. *mCherry^RNAi^*: n= 10, m= 34.85 ± 4.22. *UAS-Dap,UAS-Rbf*: n= 8, m= 18.02 ± 4.19. *p < 0.05. F. Quantification of the differentiation status of the tumours in A-B’’, measured by Elav^+^ neuronal volume as a percentage of DAPI^+^ brain volume, performed using an unpaired t-test. *mCherry^RNAi^*: n= 6, m= 14.79 ± 1.35. *UAS-Dap,UAS-Rbf*: n= 6, m= 16.76 ± 0.78. ns (not significant) p > 0.05. G-H. Representative images of brain lobes, where *mCherry^RNAi^* or *dup^RNAi^* was induced in CG cells of tumour-bearing animals from 0 h AEL using *elavQF>Qpros^RNAi^*, *nrv2-GAL4,UAS-GFP* and dissected at 120 h AEL. NBs were marked by Dpn, and glial cells by Repo. Scale bar= 50 μm. I. Quantification of glial cell number (Repo^+^, FC) in G-H. Data normalised to controls. *mCherry^RNAi^*: n= 12, m= 1 ± 0.02. Data normalised to controls. *dup^RNAi^*: n= 10, m= 0.84 ± 0.04. **p < 0.01. J. Quantification of brain size (FC) in G-H, performed using an unpaired t-test. Data normalised to controls. *mCherry^RNAi^*: n= 22, m= 1 ± 0.04. *dup^RNAi^*: n= 27, m= 0.8 ± 0.03. ***p < 0.001. K-L. Representative images of brain lobes, where *UAS-hid,rpr* was induced in CG cells of tumour-bearing animals from 96 h AEL vs control, using *elavQF>Qpros^RNAi^*, *nrv2-GAL4, UAS-GFP,GAL80^ts^,* and dissected at 144 h AEL. NBs were marked by Dpn, glial cell nuclei by Repo, and CG membranes by GFP. Scale bar= 50 μm. M. Quantification of glial cell number (Repo^+^, FC) in K-L, performed using an unpaired t-test. Data normalised to controls. *+*: n= 7, m= 1 ± 0.04. *UAS-hid,rpr*: n= 8, m= 0.28 ± 0.01. ****p < 0.0001. N. Quantification of brain size (FC) in K-L, performed using a Mann-Whitney test. Data normalised to controls. *+*: n= 7, m= 1 ± 0.04. *UAS-hid,rpr*: n= 8, m= 0.37 ± 0.04. ***p < 0.001. O-P’. Representative images of brain lobes, where *lacZ* or *shi^ts^*were induced in CG cells of tumour-bearing animals from 72 h AEL using *elavQF>Qpros^RNAi^*, *nrv2-GAL4,UAS-GFP* and dissected at 144 h AEL. NBs were marked by Dpn, glial cell nuclei by Repo, and CG membranes by GFP. Scale bar= 50 μm. Q. Quantification of glial cell number (Repo^+^, FC) in O-P, performed using an unpaired t-test. Data normalised to controls. *lacZ*: n= 11, m= 1 ± 0.07. *shi^ts^*: n= 12, m= 0.61 ± 0.04. ****p < 0.0001. N. Quantification of NB cell number (Dpn^+^, FC) in O-P, performed using an unpaired t-test. Data normalised to controls. *lacZ*: n= 11, m= 1 ± 0.03. *shi^ts^*: n= 12, m= 0.56 ± 0.03. ****p < 0.0001. O. Summary diagram – Expanded CG niche in *pros* KD-mediated tumours is generated during the Syp^+^ temporal window and is mediated by two mechanisms: 1. Gliogenesis from tumour NBs, and 2. Prolonged proliferation of CG cells.

Next, we disrupted secreted signals from the niche, as these signals have been implicated in the regulation of neurogenesis (Morante *et al*, 2013; Cheng *et al*, 2011; Bailey *et al*, 2015; Spéder & Brand, 2018; Dumstrei *et al*, 2003; Read, 2018). We inhibited secretary signals from the CG (*nrv2-GAL4*) by expressing a temperature-sensitive dominant negative form against the dynamin *shibere* (*shi^ts^*) (Figure 6 O-R, from 72 h AEL) or an RNAi against the endocytic machinery *rab11* (Figure S5 I-L, from the start of embryogenesis). We found that inhibiting the secretion of signals led to disrupted glial morphology and a significant decrease in both glial and NB numbers. We then investigated whether inhibiting the secretory machinery in the CG would have the same effect on the WT brain. Expression of *shits* from 72 h AEL resulted in a reduction in both glial cell and NB numbers (Figure S5 M-P). Together, these results indicate that CG-derived secreted factors are essential for sustaining NB number and tumour growth. To examine the role of secreted factors further, we undertook a candidate approach. A number of signalling pathways activated by secreted ligands have been implicated in glial niche-NB communication. These include PI3K pathway, activated by Jellybelly (Jeb) (Cheng *et al*, 2011), Hedgehog signalling pathway activated by Hedgehog (Hh) (Dong *et al*, 2021), FGF signalling activated via Pyramus (Pyr) and Thisbe (Ths) (Avet-Rochex *et al*, 2012), EGFR signalling activated by Spitz (Spi) (Morante *et al*, 2013) and Notch signalling activated by Delta (Dl) (Sood *et al*, 2022). However, the knockdown of these ligands using the CG drivers *NP2222-GAL4* or *nrv2-GAL4* did not significantly reduce tumour growth (Figure S5 Q).

Altogether, our study shows that dedifferentiation-induced tumours produce ectopic glial cells during the Syp^+^ temporal window, which contributes to the generation of a supportive niche. Moreover, as the brain grows during development, the glial niche expands via cell divisions, and fails to exit from the cell cycle in an ecdysone signalling-dependent manner. Promoting gliogenesis via the overexpression of *repo* or by extending the Syp^+^ temporal window is sufficient to increase tumour size. Moreover, disrupting the *de novo* generated niche (via *syp* knockdown), or the glial niche in general (via inhibition of cell cycle, endoreplication or cellular growth) diminishes tumour size, suggesting that the niche is required for tumour growth.

## Discussion

Tumour cells have been shown to exhibit characteristics of stem cells, as they are multipotent and can give rise to different cell fates with different functions (Batlle & Clevers, 2017). These cancer stem cells (CSCs) can be derived from committed and differentiated cell types, via a process called dedifferentiation. The increased plasticity of CSCs has been shown to contribute to the construction of a microenvironment which supports tumour growth and metastasis in some tissue contexts (Tammela *et al*, 2017). In the *Drosophila* CNS, while dedifferentiation has been well characterised, the interaction between tumours formed by this process and their immediate niche has not been explored. In this study, we show that NSCs that arise via dedifferentiation are capable of giving rise to CG cells which constitute their immediate niche. This process of *de novo* gliogenesis, coupled with an extended period of glial proliferation, contributes to the expansion of the CG niche, which in turn facilitates the growth and progression of the tumour (Figure 6 S).

During homeostasis, CG cells form syncytial units surrounding NB lineages and provide them with structural and metabolic support (Rujano *et al*, 2022; Spéder & Brand, 2018). Although CG numbers have been shown to increase exponentially during larval stages (Pereanu *et al*, 2005; Rujano *et al*, 2022), mitoses occur at relatively low frequencies (1%, as reported by Pereanu *et al*, 2005 and Rujano *et al*, 2021), suggesting that this mechanism alone cannot account for CG expansion. Gliogenesis via neuroglioblast progenitors has been suggested to be a likely mechanism that contributes to the expansion of the glial cell pool (Pereanu *et al*, 2005). Our G-TRACE and clonal analysis in type I WT lineages support this hypothesis as we observe glial cells which are derived from NB lineages. This suggests that WT CG expansion could be attributed to by gliogenesis from NB lineages as well as their continuous proliferation during larval stages. In this study, we have mainly focused on how the tumour CG niche forms, and the role it plays. Nonetheless, investigating the mechanisms underlying *de novo* gliogenesis in WT lineages will be an interesting area to explore in the future.

We found that the CG niche expansion in tumour-bearing animals is mediated by *de novo* gliogenesis from tumour NBs. While the glial master regulators Gcm and Repo are not required for *de novo* gliogenesis in the tumour, Repo is sufficient to drive the expansion of the glial niche, and this in turn sustains tumour growth. In follow-up studies, it would be important to identify the mechanism by which glial niche expansion promotes tumour growth, whether it is via secreted factors, cell adhesion mediators or metabolic factors.

We show that temporal patterning regulates the timing at which NBs have the capability to give rise to CG cells. During development, temporal patterning progression ensures the production of the right types and numbers of neural progeny (Maurange, 2020). Opposing temporal gradients of the RNA-binding proteins Imp and Syp regulate neuronal fate specification through a hierarchical gene regulatory network (Ren *et al*, 2017; Liu *et al*, 2015; Syed *et al*, 2017). In the context of tumorigenesis, Imp^+^ NBs have been shown to be highly proliferative and are capable of conferring the malignant competency of tumours (Narbonne-Reveau *et al*, 2016; Genovese *et al*, 2019; van den Ameele & Brand, 2019). Our study shows the more differentiated Syp^+^ NBs also plays an important role in regulating tumour growth. Syp is responsible for the formation CG cells, and its expression is necessary and sufficient to promote tumour growth. Importantly, extending the Syp^+^ temporal window promotes the expansion of the CG niche, and in turn promotes tumour growth. The role of Syp in regulating glial cell formation has been reported in type II NBs (Syed *et al*, 2017). Similar to mammalian neurogenesis, these NBs produce glial cells following a neurogenic phase (Jacobson, 1991). Mammalian neurogenesis has also been shown to be regulated by temporal patterning. For example, the Chick Ovalbumin Upstream Promoter-Transcription Factors (COUP-TFI and II) is the orthologue of *Drosophila* temporal regulator Seven-Up (Svp), previously implicated in promoting the transition between early and late temporal cell fates (Maurange *et al*, 2008; Syed *et al*, 2017). COUP-TFs are expressed early in neural progenitors in the developing mouse brain, where they are required for the switch between neuronal and glial production (Naka *et al*, 2008). Our data, together with these studies, suggest that temporal progression regulates the production of glial cells in neural development and brain tumour formation.

Another contributor to the overall expansion of the glial niche is via prolonging the overall period of CG proliferation, mediated via ecdysone signalling. It is known that ecdysone signalling is dependent on the level of systemic steroid hormone ecdysone, which peaks at late larval stages to stop larval growth (Riddiford, 1993). It has been shown that wing disc tumours can cause developmental delay via a reduction in the overall ecdysone titre (Sehnal & Bryant, 1993; Colombani *et al*, 2012). Here, we show that brain tumours also enlist similar mechanisms to reduce systemic ecdysone levels.

CG has been shown to be required for NB proliferation and survival (Dong *et al*, 2021; Read, 2018) and resistance to environmental stresses (Cheng *et al*, 2011; Bailey *et al*, 2015), as well as to influence NBs’ decision between quiescence and reactivation (Sousa-Nunes *et al*, 2011; Chell & Brand, 2010). Furthermore, impaired CG growth and proliferation can lead to neuronal apoptosis (Spéder & Brand, 2018; Rujano *et al*, 2022). In the context of tumour growth, we show that *de novo* gliogenesis, as well as overall niche integrity, are important for sustaining tumour growth. This parallels other cancer contexts where tumour progression relies on the niche and genetic perturbation of the niche results in inhibited tumour potential (Tammela *et al*, 2017). Overall, our study shows that tumour remodels and is capable of generating its own microenvironment to support its malignant growth. Targeting the niche, thus, could be a potential avenue exploited in the clinic to inhibit the tumour stem cell-like program in brain cancers.

## Materials and Methods

### Fly husbandry and strains

*Drosophila melanogaster* stocks were raised on standard medium at room temperature. Mated animals were left at 25 °C to lay eggs for 24 h, the crosses were then moved to 29 °C unless otherwise stated and dissected at late third instar. Control and tumour-bearing animals were reared under the same conditions and brains were dissected at the 120 h AEL time point for direct comparison. For experiments with only tumour brains, dissections were performed at either 120 or 144 h AEL, as stated in the figure legends. The fly strains used in this study are listed in Supplementary Table 1.

### Clone induction

WT (*FRT82B*) and *pros* mutant (*FRT82B;pros^17^*) clones were induced using the MARCM system: *UAS-nGFP,hsFLP;FRT82B, tub-GAL80; tub-GAL4*. Clones were induced via heat-shock at 48 h AEL for 15 min (WT) or 8 min (*pros^17^*) at 37 °C. Animals were raised at 25°C and dissected at 120 h AEL (larvae) or 24 h after pupa formation (pupae).

### Immunostaining

Brains were dissected in PBS, fixed for 20 minutes in 4% formaldehyde (Sigma-Aldrich, #F8775) in PBS and washed in 0.5% PBST (PBS + 0.5% TritonX-100 (Sigma-Aldrich, #T8787)). Brains were incubated with primary and secondary antibodies in 0.5% PBST overnight at 4°C. Samples were mounted in 80% glycerol in PBS for image acquisition. The primary antibodies used were: rat anti-Dpn (1:200; Abcam, 195172), mouse anti-Repo (1:50; DSHB, 8D12), chick anti-GFP (1:2000; Abcam, 13970), mouse anti-Elav (1:100; DSHB, 9F8A9), rabbit anti-Ase (a gift from Lily Jan and Yuh Nung Jan), mouse EcR-B1 (1:50; DHSB, AD4.4), chicken anti-β-gal (1:1000; Abcam, 9361). Secondary antibodies (Molecular Probes) were used at 1:500. DAPI (Molecular Probes) was used at 1:10000.

### EdU labelling

EdU labelling was used to detect actively dividing NBs (Figure 1 D-E’, G, Figure 6 A’’-B’’, E). Larval brains (120 h AEL) were incubated in 10uM EdU/PBS for 15 minutes, followed by fixation. Brains were then stained against anti-Dpn and anti-Repo to label NBs and glial cells, respectively, before EdU detection using the Click-iT EdU Cell Proliferation Kit (Invitrogen, C10340).

EdU feeding was performed to detect the number of dividing glial cells over a 24 h time window (Figure 2 B-E). Larvae staged at 72 h AEL were fed with standard food supplemented with 100ug/mL EdU. Brains were dissected 22 h later and fixed, followed by Repo staining and EdU detection as indicated above.

### 20E feeding

20-Hydroxyecdysone (20E, Sigma, H5142-5MG) was dissolved in 1 mL of 100% ethanol (Sigma, 459836) to make a 5 mg/mL stock solution (stored at −20 °C). 20E stock solution or 100% ethanol was mixed into standard food (final concentration of 0.15 mg/mL) (Nogueira Alves *et al*, 2022). The food supplemented with 20E or ethanol was air dried at room temperature overnight before use. Larvae, staged at 72 h AEL, were fed on the food at 96 h AEL and dissected at 120 h AEL.

### Pupariation

For pupariation experiment in Figure S1 A, control and tumour-bearing larvae (30 each) were placed in fresh food vials at 72 h AEL (4 replicates per condition). The number of larvae that pupated was scored on consecutive days.

For pupariation experiment upon 20E feeding in Figure 3 F, tumour-bearing animals were staged at the L2-L3 moulting stage (∼72 h AEL). Larvae were moved to standard food supplemented with ethanol or 20E in ethanol (as described above) and the number of larvae that pupated was scored on consecutive days.

### Molecular cloning

The *QUAS-pros^RNAi^* strain was generated in this study. The *pros^RNAi^* construct was amplified from genetic DNA of the BDSC42538 TRiP line, using the oligos (Sigma-Aldrich) listed in Table 9. The resulting PCR product was digested with EcoRI HF and NheI HF. The digested fragment was cloned into a pQUAS-WALIUM20 vector (DGRC, Stock 1474), following the manufacturer’s protocol. The plasmid was transferred into SURE Supercompetent cells (Agilent Technologies, 200152). The *QUAS-pros^RNAi^* transgene was injected into flies carrying the attP-3B (BDSC9725) and attP-9A (BDSC9752) docking sites by BestGene Inc.

### Quantitative real-time PCR (RT-qPCR)

15 brain-ring gland complexes (BRGC) were dissected from control (*elavQF>+*) and tumour (*elavQF>Qpros^RNAi^*) larvae at 120 h AEL (3 replicates per genotype). RNA was extracted using the Direct-zol RNA MicroPrep Kit (ZYMO Research, #R2060) and reverse-transcribed using the ProtoScript II First Strand cDNA Synthesis Kit (NEB, #E6560). Synthesised cDNAs were then diluted 1:10. Then Fast SYBR Green Master Mix (Thermo Fisher Scientific, #4385612) was used for qPCR by StepOnePlus qPCR machine (Applied Biosystems). The ΔΔCT method was used to calculate fold changes, and transcript levels were normalized to *rpl42* expression. Statistical significance was tested with two independent unpaired student t-tests. qPCR primers are listed in Table 9.

### Image acquisition and processing

Images were collected on an Olympus FV3000, using 2 μm-spaced Z stacks. Images were processed with https://imagej.net/software/fiji/. The ventral side of the CB, where type I NBs reside, is always shown as the representative image. Image brightness adjustments were applied equally to control and experimental conditions to allow for better visualisation when required.

### Image quantification and analysis

#### Brain size and NB volume measurements

Brain size and NB volume were used interchangeably as proxies of tumour size, as *elav-QF* and *dnab-GAL4* -driven *pros^RNAi^* tumours occupied most of the brain. Brain size was calculated by measuring the brain area of maximum projections of whole brain images. NB (Dpn^+^) volume was measured from three-dimensional reconstructions of confocal Z-stacks with Volocity software (Improvision) or Imaris (Bitplane). In the WT brain, the number of type I NBs was counted manually (Figure S3 I-K, Figure S4 G-I and K-M, Figure S5 A-B, D and M-N, P). Data was normalised to their respective controls.

#### Glial cell number and glial volume measurements

Glial cell number or glial volume (from Repo^+^ cells) were measured from three-dimensional reconstructions of confocal Z-stacks with Volocity software (Improvision) or Imaris (Bitplane). Glial cell number was also automatically measured with Fĳi’s plug-in “DeadEasy Larval Glia” (Forero *et al*, 2012). No significant differences were observed when glial cell number was measured compared to glial volume (see graph below). Data was normalised to their respective controls. CG nuclear size was calculated by measuring the cell area of Repo^+^ and *NP2222-GAL4>UAS-GFP* positive cells.

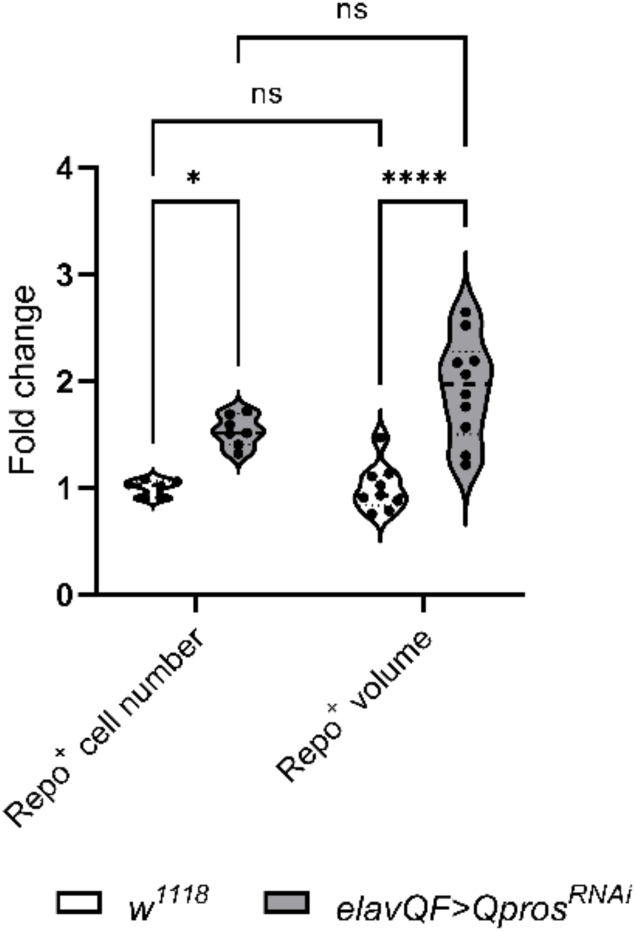

#### EdU index

EdU is incorporated into cells during the S phase of the cell cycle. The EdU index is the percentage of cells marked by EdU as a proportion of the total number of cells. In Figure 1 G and Figure 6 E, the NB EdU index was measured with Volocity as follows: EdU^+^ Dpn^+^ volume/total Dpn^+^ volume. In Figure 2 D, number of Edu^+^ Repo^+^ cells was counted manually. In Figure 2 E, the number in D was divided by the total number of Repo^+^ cells to show the EdU index.

#### FUCCI

Fly FUCCI was expressed using the GAL4-UAS system in CG, where cells in G2 or M phase are marked with both nuclear ECFP and Venus, S phase with Venus only, and G1 phase with ECFP only. The number of cells in each cell cycle phase was automatically calculated with a formula and a glial cell counting macro (Computer Code Macro 1; adapted from (Forero *et al*, 2012)). Specifically, the total number of marked cells (n^Total^), cells marked by ECFP only (n^ECFP+^), and cells marked by Venus only (n^Venus+^) were automatically counted. The number of cells in each phase was calculated as below:

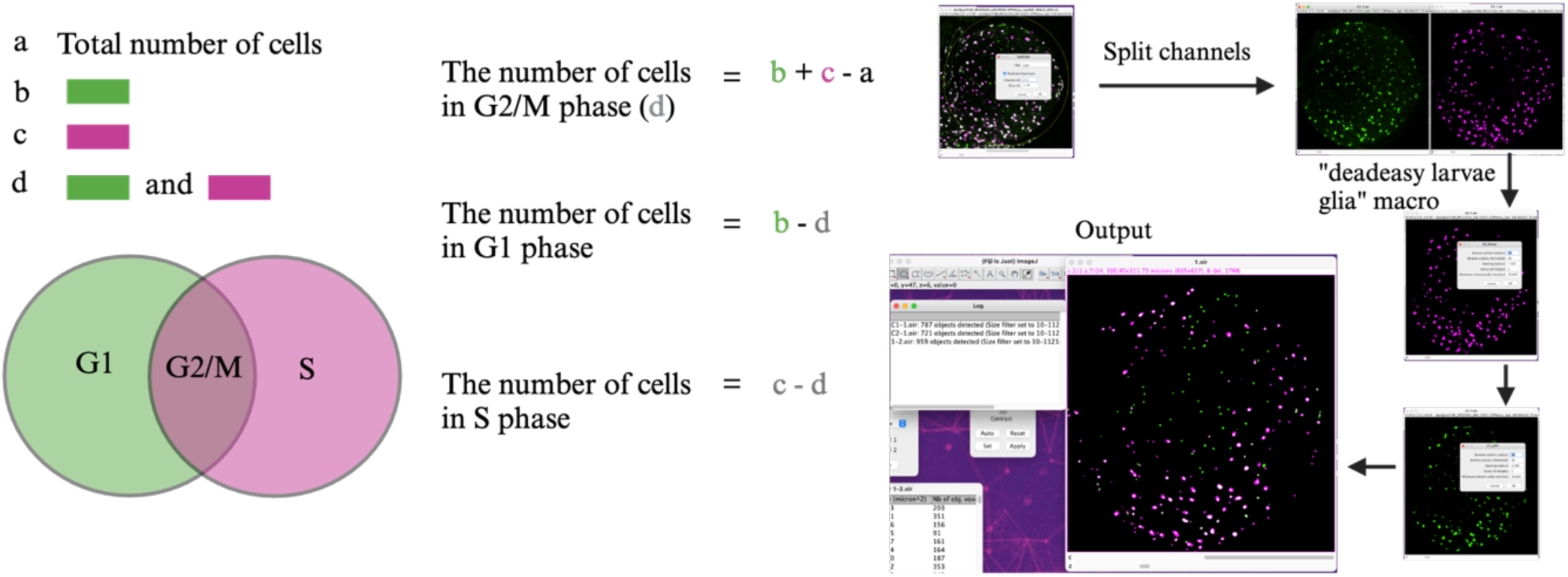

Compared with manual counting, automatic counting excluded weak ECFP^+^ cells, however, this did not affect our conclusion. Chi-square tests were used to determine the difference in CG cell cycle distribution between control and tumour-bearing larvae.

#### CTCF measurement

In Figure 3 B-D, the intensity of EcR-B1 in CG was measured in Fĳi using the following formula: CTCF (corrected total cell fluorescence) = Integrated density – (Area of CG cell x mean fluorescence of background readings) (McCloy *et al*, 2014). CG cells were marked with *wrapper-GAL4>UAS-RFP.nls* expression.

### Statistical analysis

All data are plotted in violin plots, which depict the distribution of the data points using density curves. The three lines within the violin plot indicate the quartile positions. All statistical analyses were conducted using GraphPad Prism 9.0 (©GraphPad 593 software Inc.). For experiments with two genotypes or conditions, two-tailed unpaired student’s t-tests were used to test for significant differences. The Welch’s correction was applied in cases of unequal variances, and the Mann-Whitney U test was used in the cases of violated normality. For experiments with more than two genotypes or conditions, significant differences between specific genotypes were tested using a one-way ANOVA and a subsequent Šidák post-hoc test. A Brown-Forsythe correction was applied in cases of unequal variances, and in the cases of violated normality, the Kruskal-Wallis test was used. For all histograms, error bars represent SEM. p and adjusted-p values are reported as follows: p>0.05, ns (not significant); p<0.05, *; p<0.01, **; p<0.001, ***; p<0.0001, ****.

## Acknowledgements

We are grateful to Angela Giangrande, Cedric Maurange, Lily Jan and Yuh Nung Jan for generous sharing of fly stocks and antibodies. We would like to thank Michael Zavortink for expert assistance with molecular biology. We would also like to thank Khanh Nguyen, Kieran Harvey and Sarah Russell for critical reading of the manuscript, and members of the Cheng lab for fruitful discussions. We also thank the Bloomington Drosophila Stock Centre, Vienna Drosophila Resource Centre, BestGene Inc and Developmental Studies Hybridoma Bank for fly stocks, embryo injections and antibodies. We would like to also thank OZDros for *Drosophila* quarantine and the Peter MacCallum Cancer Institute CAHM facility for microscopy assistance. EAO was supported by a Melbourne Research Scholarship from The University of Melbourne. LYC’s laboratory is supported by funding from the NHMRC Ideas Grant (APP2011289), and the Peter MacCallum Cancer Foundation.

## Disclosure and competing interests statement

The authors declare no competing interests.

## Notes

### Competing Interest Statement

The authors have declared no competing interest.

