## supplemental file for "Tumour-derived gliogenesis sustains dedifferentiation-dependent tumour growth in the *Drosophila* CNS"

#### Supplemental figures and tables

##### Figure S1

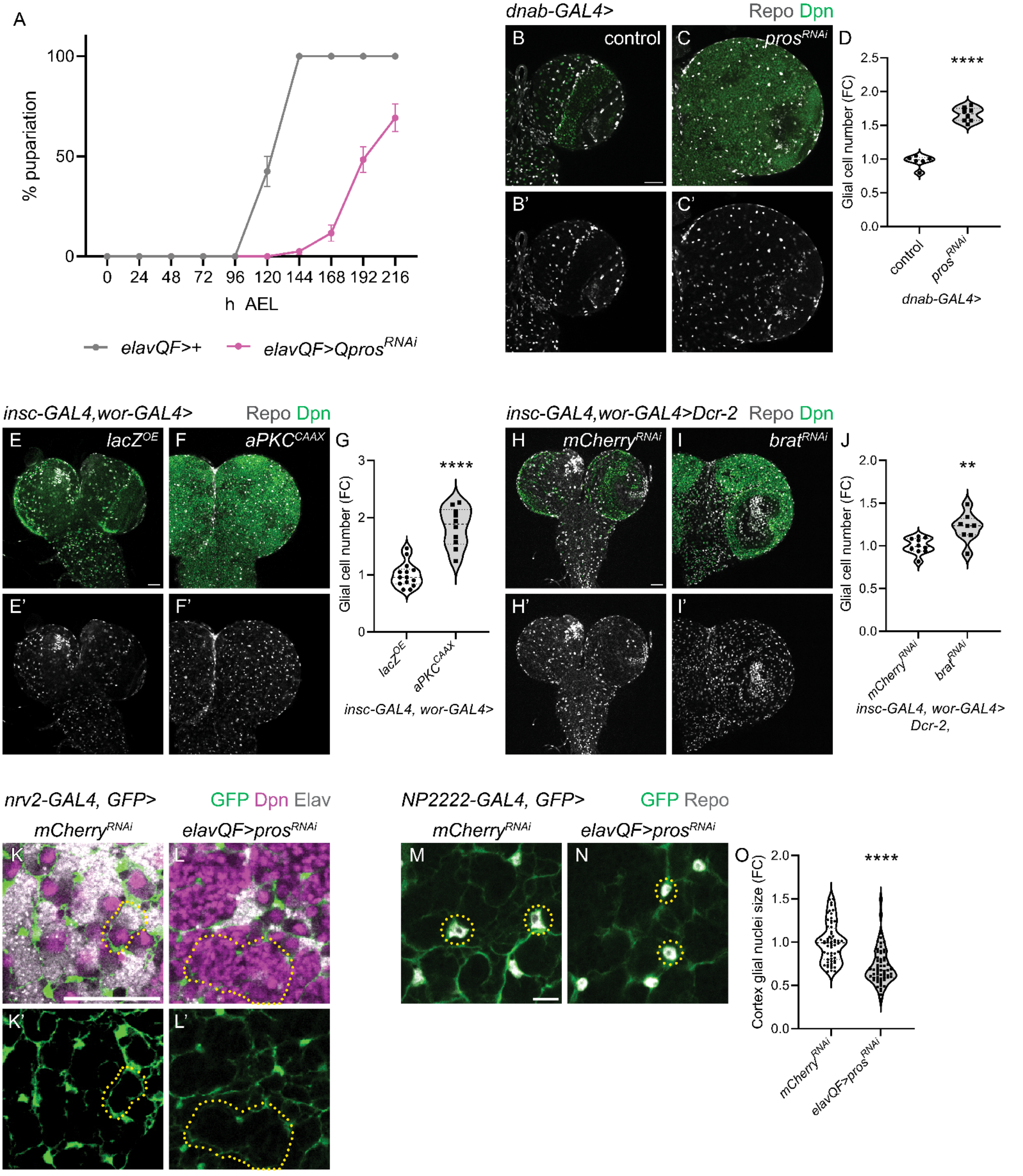

**Figure S1. Characterisation of the developmental delay, glial cell number change in other brain tumour types and glial niche alterations.**

A. *elavQF>Qpros$^{RNAi}$* tumour-bearing animals exhibit delayed pupariation compared to *elavQF*>+ control. Statistical analysis performed using a two-way ANOVA followed by Šídák's multiple comparisons post-hoc test. See Table 2 for n, mean, SEM, and adjusted p-values.

B-C'. Representative images of brain lobes where *dnab-GAL4>pros$^{RNAi}$* exhibit increased glial cell number compared to control *dnab-GAL4*>+. Glial cells are marked by Repo, NBs are marked by Dpn. Scale bar= 50 µm.

D. Quantification of glial cell number in B-C', performed using an unpaired t-test. Data normalised to controls. Control: n= 6, m= 0.97 ± 0.04. *pros$^{RNAi}$*: n= 8, m= 1.67 ± 0.04. ****p < 0.0001.

E-F'. Representative images of brain lobes where *insc-GAL4,wor-GAL4>UAS-Dcr2, UAS-aPKC$^{CAAX}$* exhibit increased glial cell number compared to *Insc-GAL4,wor-GAL4>UAS-lacZ*. Glial cells are marked by Repo, NBs are marked by Dpn. Scale bar= 50 µm.

G. Quantification of glial cell number in E-F', performed using an unpaired t-test. Data normalised to controls. *lacZ$^{OE}$*: n= 14, m= 1 ± 0.06. *aPKC$^{CAAX}$*: n= 10, m= 1.83 ± 0.11. ****p < 0.0001.

H-I'. Representative images of CNSs, *insc-GAL4,wor-GAL4>UAS-Dcr-2,brat$^{RNAi}$* exhibit increased glial cell number compared to *insc-GAL4,wor-GAL4>UAS-Dcr-2,mCherry$^{RNAi}$*. Glial cells are marked by Repo, NBs are marked by Dpn. Scale bar= 50 µm.

J. Quantification of glial cell number in H-I', performed using an unpaired t-test. Data normalised to controls. *mCherry$^{RNAi}$*: n= 10, m= 1 ± 0.03. *brat$^{RNAi}$*: n= 8, m= 1.22 ± 0.06. **p < 0.01.

K-L'. Representative images of zoomed-in brain lobes, where CG membranes marked by *nrv2-GAL4,UAS-GFP* in *elavQF>pros$^{RNAi}$* enclose multiple NBs compared to *elavQF>mCherry$^{RNAi}$*. Glial membrane is marked by GFP, NBs are marked by Dpn, and neurons are marked by Elav. Glial chambers enwrapping NB lineages are outlined. Scale bar= 50 µm.

M-N. Representative images of zoomed-in brain lobes, where CG cells marked by *NP2222-GAL4,UAS-GFP* and Repo exhibit smaller nuclear size in *elavQF>pros$^{RNAi}$* compared to *elavQF>mCherry$^{RNAi}$*. The CG nuclei are outlined in yellow. Scale bar= 10 µm.

O. Quantification of glial nuclear size in M-N, performed using a Mann-Whitney test. Data normalised to controls. *mCherry$^{RNAi}$*: n= 60, m= 1 ± 0.03. *elavQF>pros$^{RNAi}$*: n= 60, m= 0.72 ± 0.03. ****p < 0.0001.

### Figure S2

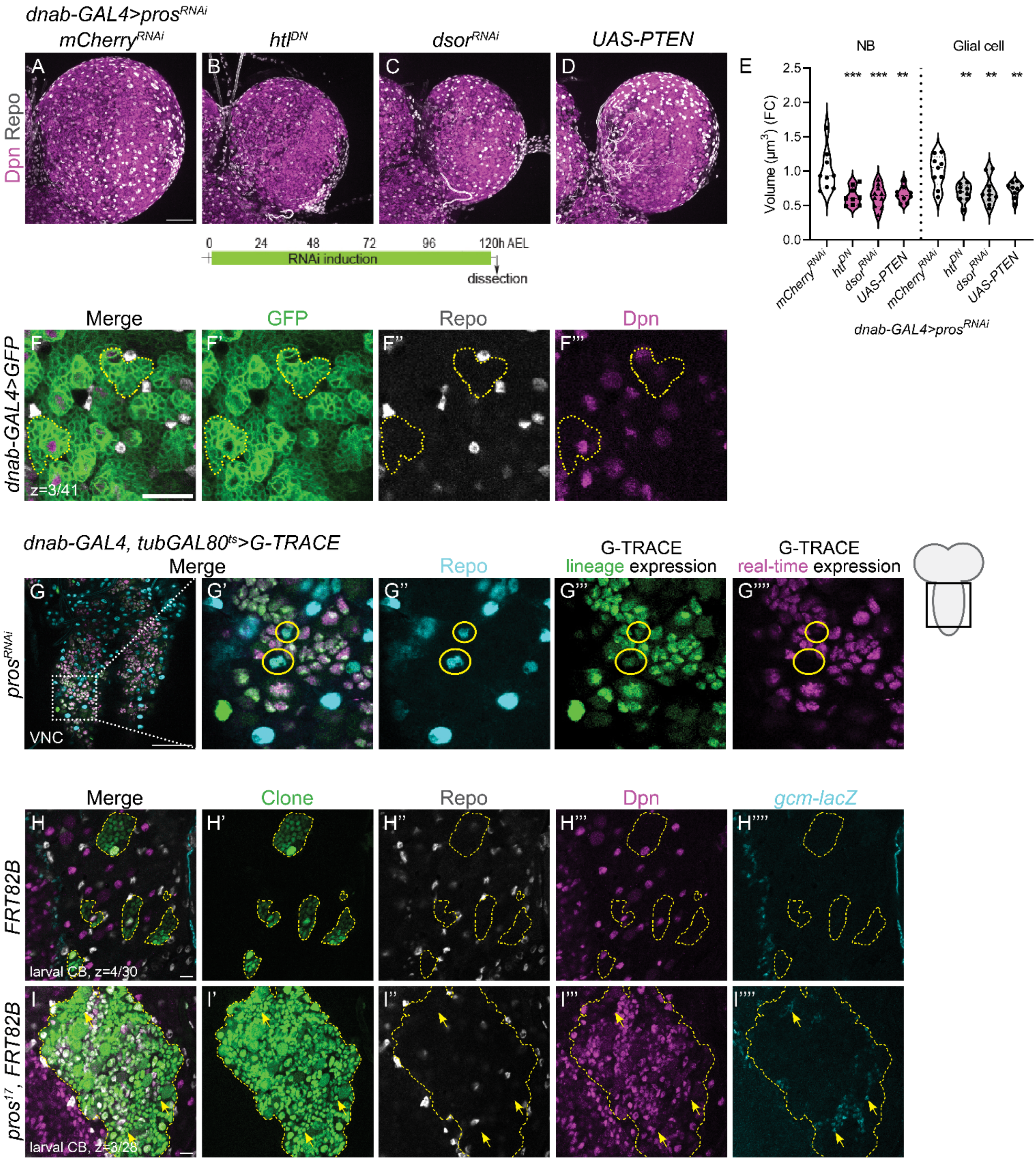

dnab-GAL4>pros^RNAi
mCherry^RNAi
htl^DN
dsor^RNAi
UAS-PTEN
Dpn Repo
A
B
C
D
E
NB
Glial cell
Volume (µm³) (FC)
2.5
2.0
1.5
1.0
0.5
0.0
*** *** **
** ** **
mCherry^RNAi htl^DN dsor^RNAi UAS-PTEN mCherry^RNAi htl^DN dsor^RNAi UAS-PTEN
dnab-GAL4>pros^RNAi
0 24 48 72 96 120h AEL
RNAi induction
dissection
Merge
GFP
Repo
Dpn
dnab-GAL4>GFP
F F' F'' F'''
z=3/41
dnab-GAL4, tubGAL80ts>G-TRACE
Merge
Repo
G-TRACE lineage expression
G-TRACE real-time expression
pros^RNAi
G G' G'' G''' G''''
VNC
Merge
Clone
Repo
Dpn
gcm-lacZ
FRT82B
H H' H'' H''' H''''
larval CB, z=4/30
pros^17, FRT82B
I I' I'' I''' I''''
larval CB, z=3/28

**Figure S2. Tumours express the glial cell fate regulator *gcm* and generate glial cells**

A-D. Representative images of brain lobes (maximum projections) at 120 h AEL where $htl^{DN}$, $dsor^{RNAi}$ and *UAS-PTEN* expression in *dnab-GAL4>*$pros^{RNAi}$ from 0 h AEL caused reduced tumour size and glial cell number compared to the expression of $mCherry^{RNAi}$. Glial cells are marked by Repo and NB by Dpn. Scale bar= 50 µm.

E. Quantification of NB and glial cell volume (FC) in A-D, performed using ordinary one-way ANOVAs. Data normalised to controls. NB volume – $mCherry^{RNAi}$: n= 9, m=1 ± 0.1. $htl^{DN}$: n= 9, m= 0.63 ± 0.4. $dsor^{RNAi}$: n= 10, m= 0.63 ± 0.05. *UAS-PTEN*: n= 9, m= 0.66 ± 0.04. Glial cell volume – $mCherry^{RNAi}$: n= 9, m=1 ± 0.08. $htl^{DN}$: n= 9, m= 0.68 ± 0.4. $dsor^{RNAi}$: n= 10, m= 0.7 ± 0.06. *UAS-PTEN*: n= 9, m= 0.73 ± 0.04. $^{**}p < 0.01$, $^{***}p < 0.001$.

F-F'''. Representative images showing that *dnab-GAL4* lineages (marked by *UAS-GFP*) contain a single NB ($Dpn^+$) and many $Dpn^-$ progeny cells, but not glial cells ($Repo^+$). Scale bar= 20 µm.

G-G''''. Representative images showing tumour lineages (*dnab-GAL4>*$pros^{RNAi}$) within the VNC expressing G-TRACE. Real-time expression is visualised in magenta, lineage expression in green and glial cells in cyan. G'-G''' are zoomed in images from G. Glial cells with G-TRACE lineage expression, but no real-time expression, are circled. Scale bar= 50 µm.

H-I. Representative images showing that *gcm-lacZ* is expressed within $pros^{17}$ MARCM clones, but not in WT (*FRT82B*) clones. Clones showed are in the ventral side of larval CBs. Dotted lines outline the clones and yellow arrows point to *gcm-lacZ*-expressing cells. *gcm-lacZ* expression did not collocalise with glial marker Repo or NB marker Dpn. Scale bar= 10 µm.

### Figure S3

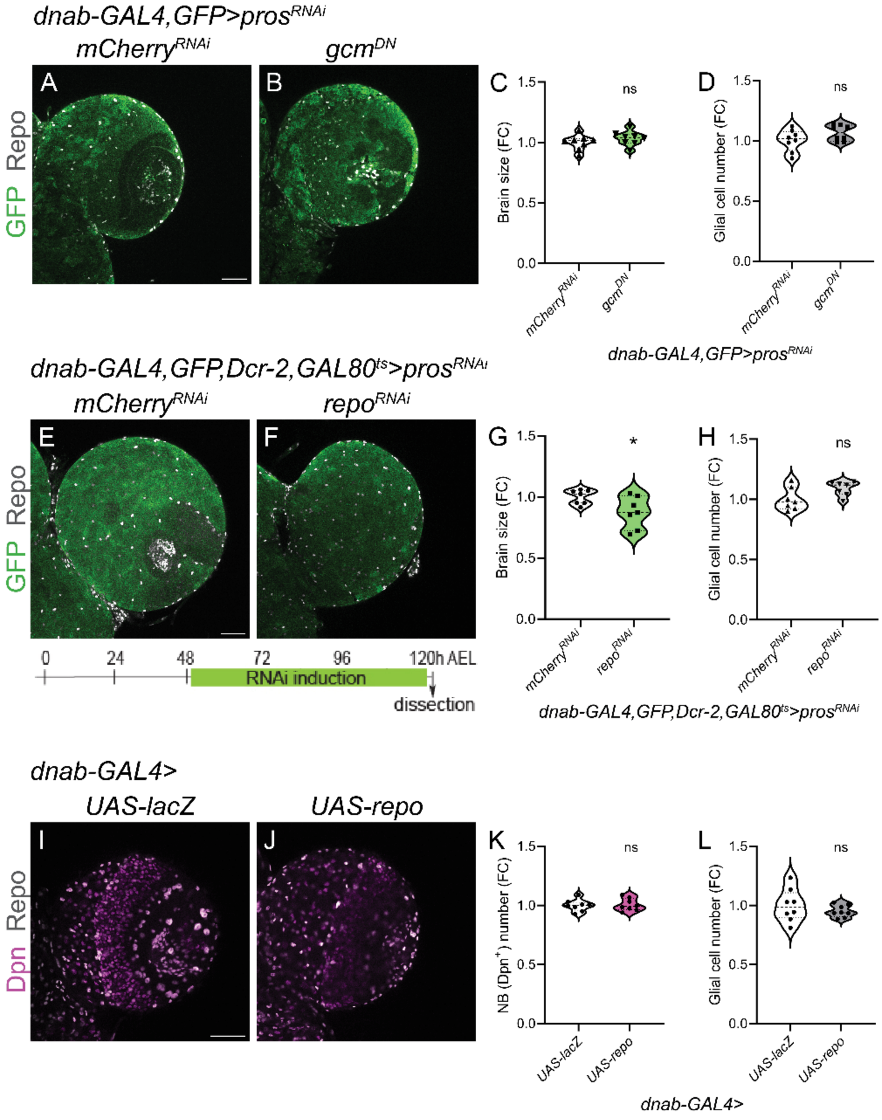

dnab-GAL4,GFP>pros^RNAi
mCherry^RNAi
gcm^DN
A
B
GFP Repo
C
D
Brain size (FC)
Glial cell number (FC)
ns
ns
dnab-GAL4,GFP>pros^RNAi
dnab-GAL4,GFP,Dcr-2,GAL80^ts>pros^RNAi
mCherry^RNAi
repo^RNAi
E
F
G
H
*
ns
0
24
48
72
96
120h AEL
RNAi induction
dissection
dnab-GAL4,GFP,Dcr-2,GAL80^ts>pros^RNAi
dnab-GAL4>
UAS-lacZ
UAS-repo
I
J
Dpn Repo
K
L
NB (Dpn^+) number (FC)
Glial cell number (FC)
ns
ns
dnab-GAL4>

**Figure S3. Characterisation of the role of glial cell fate genes in regulating gliogenesis from type I and tumour NBs**

A-B. Representative images of brain lobes where *gcm*$^{DN}$ is expressed in *dnab-GAL4,GFP>pros*$^{RNAi}$ tumours from 0 h AEL, where no reduction in brain size or glial number was detected compared to the *mCherry*$^{RNAi}$ control. Glial cells are marked with Repo. Scale bar= 50 µm.

C. Quantification of brain size (FC) in A-B, performed using a Kruskal-Wallis test. Data normalised to controls. The *mCherry*$^{RNAi}$ control is the same as in Figure 5 K and N. *mCherry*$^{RNAi}$: n= 8, m=1 ± 0.2. *gcm*$^{DN}$: n= 8, m= 1.03 ± 0.02. ns (not significant) $p > 0.05$.

D. Quantification of glial cell number (FC) in A-B, performed using an ordinary one-way ANOVA. The *mCherry*$^{RNAi}$ control is the same as in Figure 5 K and M. *mCherry*$^{RNAi}$: n= 8, m= 1 ± 0.03. *gcm*$^{DN}$: n= 8, m= 1.07 ± 0.02. ns (not significant) $p > 0.05$.

E-F. Representative images of brain lobes where *repo*$^{RNAi}$ is expressed in *dnab-GAL4,GFP, Dcr-2, GAL80*$^{ts}$*>pros*$^{RNAi}$ from 48 h AEL, and a mild reduction in brain size, but no significant reduction in glial number (based on Repo staining) was detected compared to the *mCherry*$^{RNAi}$ control. Scale bar= 50 µm.

G. Quantification of brain size (FC) in E-F, performed using an unpaired t-test. Data normalised to controls. *mCherry*$^{RNAi}$: n= 7, m=1 ± 0.02. *repo*$^{RNAi}$: n= 7, m= 0.87 ± 0.05. $*p < 0.05$.

H. Quantification of glial cell number (FC) in E-F, performed using an unpaired t-test. Data normalised to controls. *mCherry*$^{RNAi}$: n= 7, m=1 ± 0.04. *repo*$^{RNAi}$: n= 7, m= 1.09 ± 0.02. ns (not significant) $p > 0.05$.

I-J. Representative images of brain lobes where *UAS-repo* expression in *dnab-GAL4* lineage from 0 h AEL causes no reduction in NB (Dpn$^{+}$) or glial (Repo$^{+}$) number, compared to the expression of *mCherry*$^{RNAi}$. Scale bar= 50 µm.

K. Quantification of NB number (Dpn$^{+}$, FC) in I-J, performed using an ordinary one-way ANOVA. The *UAS-lacZ* control is the same as in Figure S4 K and M. Data normalised to controls. *UAS-lacZ*: n= 8, m=1 ± 0.02. *UAS-repo*: n= 8, m= 1.01 ± 0.01. ns (not significant) $p > 0.05$.

L. Quantification of glial cell number (Repo+, FC) in I-J, performed using an ordinary one-way ANOVA. The *UAS-lacZ* control is the same as in Figure S4 K and N. Data normalised to controls. *UAS-lacZ*: n= 8, m=1 ± 0.05. *UAS-repo*: n= 8, m= 0.95 ± 0.02. ns (not significant) $p > 0.05$.

### Figure S4

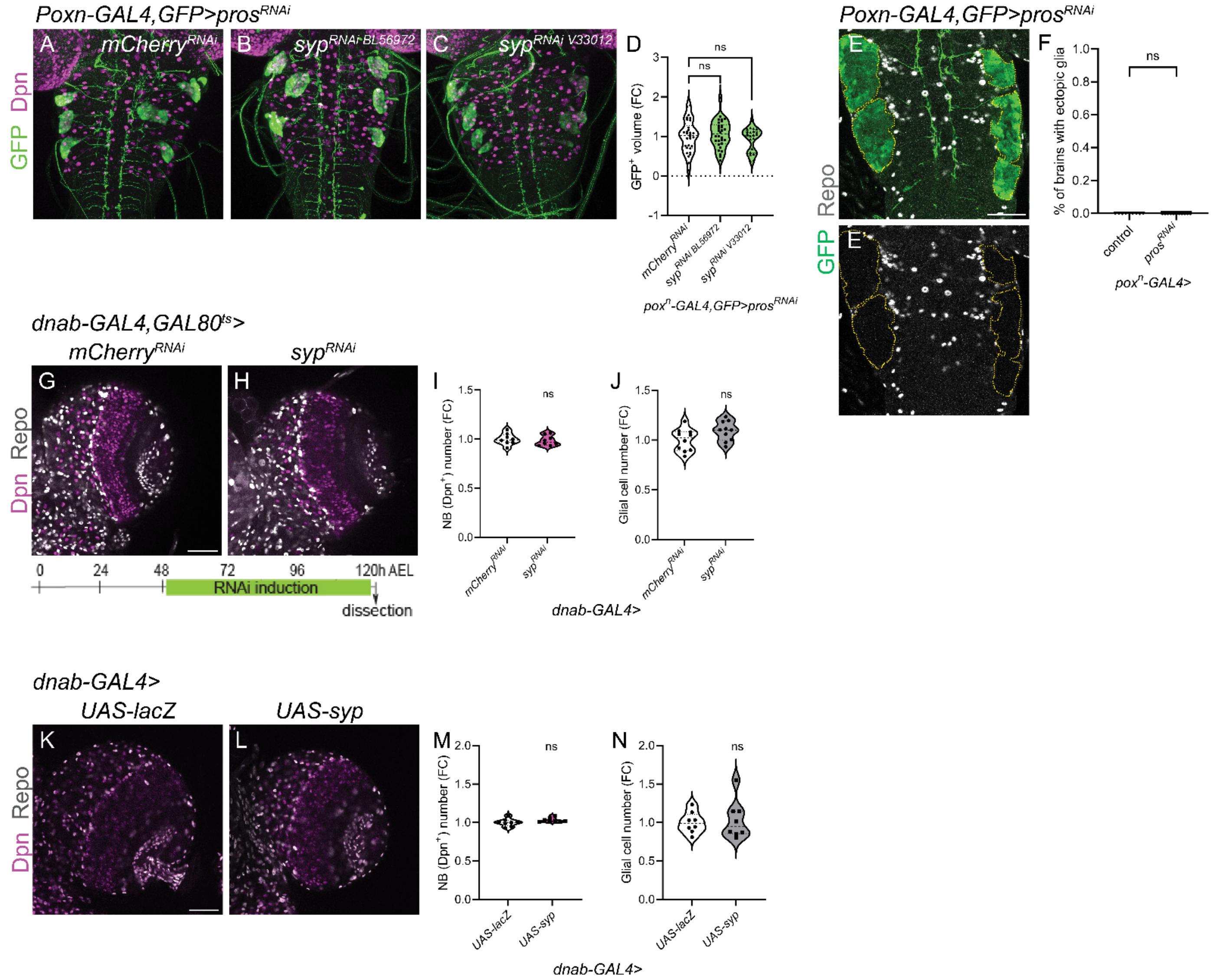

Poxn-GAL4,GFP>pros^RNAi
A mCherry^RNAi
B syp^RNAi BL56972
C syp^RNAi V33012
GFP Dpn
D
GFP+ volume (FC)
ns
ns
mCherry^RNAi
syp^RNAi BL56972
syp^RNAi V33012
pox^n-GAL4,GFP>pros^RNAi
Poxn-GAL4,GFP>pros^RNAi
E
E'
GFP Repo
F
% of brains with ectopic glia
ns
control
pros^RNAi
pox^n-GAL4>
dnab-GAL4,GAL80^ts>
mCherry^RNAi
syp^RNAi
G
H
Dpn Repo
0 24 48 72 96 120h AEL
RNAi induction
dissection
I
NB (Dpn+) number (FC)
ns
J
Glial cell number (FC)
ns
dnab-GAL4>
dnab-GAL4>
UAS-lacZ
UAS-syp
K
L
Dpn Repo
M
N
ns
ns
dnab-GAL4>

**Figure S4. Syp is not involved in NB-derived gliogenesis in wild-type nor *pros$^{RNAi}$* tumours induced in Poxn lineages.**

A-C. Representative images of VNCs where *syp$^{RNAi}$* expressed in *pox$^{n}$-GFP>pros$^{RNAi}$* lineages from 0 h AEL did not significantly alter tumour size compared to the *mCherry$^{RNAi}$* control. Scale bar= 50 µm.

D. Quantification of GFP tumour volume in A-C, performed using a Kruskal-Wallis test. Data normalised to controls. *mCherry$^{RNAi}$*: n= 30, m=1 ± 0.07. *syp$^{RNAi\ BL56972}$*: n= 30, m= 1.06 ± 0.06. *syp$^{RNAi\ V33012}$*: n= 24, m= 0.92 ± 0.06. ns (not significant) $p > 0.05$.

E-E'. Representative images of VNCs showing that no glial cells (Repo$^{+}$) were found in *Poxn*-GFP>*pros$^{RNAi}$* lineages. E' is the Repo$^{+}$ channel of the merged image in E. Scale bar= 50 µm.

F. Quantification showing no glial cells (Repo$^{+}$) are recovered in control nor *pros$^{RNAi}$ Poxn-GAL4, GFP* lineages. Analysis performed using a Mann-Whitney test. *mCherry$^{RNAi}$*: n= 9, m=0 ± 0. *pros$^{RNAi}$*: n= 0, m= 0 ± 0. ns (not significant) $p > 0.05$.

G-H. Representative images of brain lobes, where *mCherry$^{RNAi}$* or *syp$^{RNAi}$* were induced from 48 h AEL using *GAL80$^{ts}$;dnab-GAL4* and dissected at 120 h AEL. NBs were marked by Dpn, and glial cells by Repo. Scale bar= 50 µm.

I. Quantification of NB volume (Dpn$^{+}$, FC) in G-H, performed using an unpaired t-test. Data normalised to controls. *mCherry$^{RNAi}$*: n= 8, m= 1 ± 0.02. *syp$^{RNAi}$*: n= 8, m= 0.99 ± 0.02. ns (not significant) $p > 0.05$.

J. Quantification of glial cell number (Repo$^{+}$, FC) in G-H, performed using an unpaired t-test. Data normalised to controls. *mCherry$^{RNAi}$*: n= 10, m= 1 ± 0.04. *syp$^{RNAi}$*: n= 10, m= 1.09 ± 0.03. ns (not significant) $p > 0.05$.

K-L. Representative images of brain lobes, where *UAS-lacZ* or *UAS-syp* were induced from 0 h AEL using *dnab-GAL4* and dissected at 120 h AEL. NBs were marked by Dpn, and glial cells by Repo. Scale bar= 50 µm.

M. Quantification of NB volume (Dpn$^{+}$, FC) in K-L, performed using an ordinary one-way ANOVA. The *UAS-lacZ* control is the same as in Figure S3 I and K. Data normalised to controls. *UAS-lacZ*: n= 8, m=1 ± 0.02. *UAS-syp*: n= 8, m= 1.03 ± 0.01. ns (not significant) $p > 0.05$.

N. Quantification of glial cell number (Repo$^{+}$, FC) in K-L, performed using an ordinary one-way ANOVA. The *UAS-lacZ* control is the same as in Figure S3 I and L. Data normalised to controls. *UAS-lacZ*: n= 8, m=1 ± 0.05. *UAS-syp*: n= 8, m= 1.03 ± 0.09. ns (not significant) $p > 0.05$.

#### Figure S5

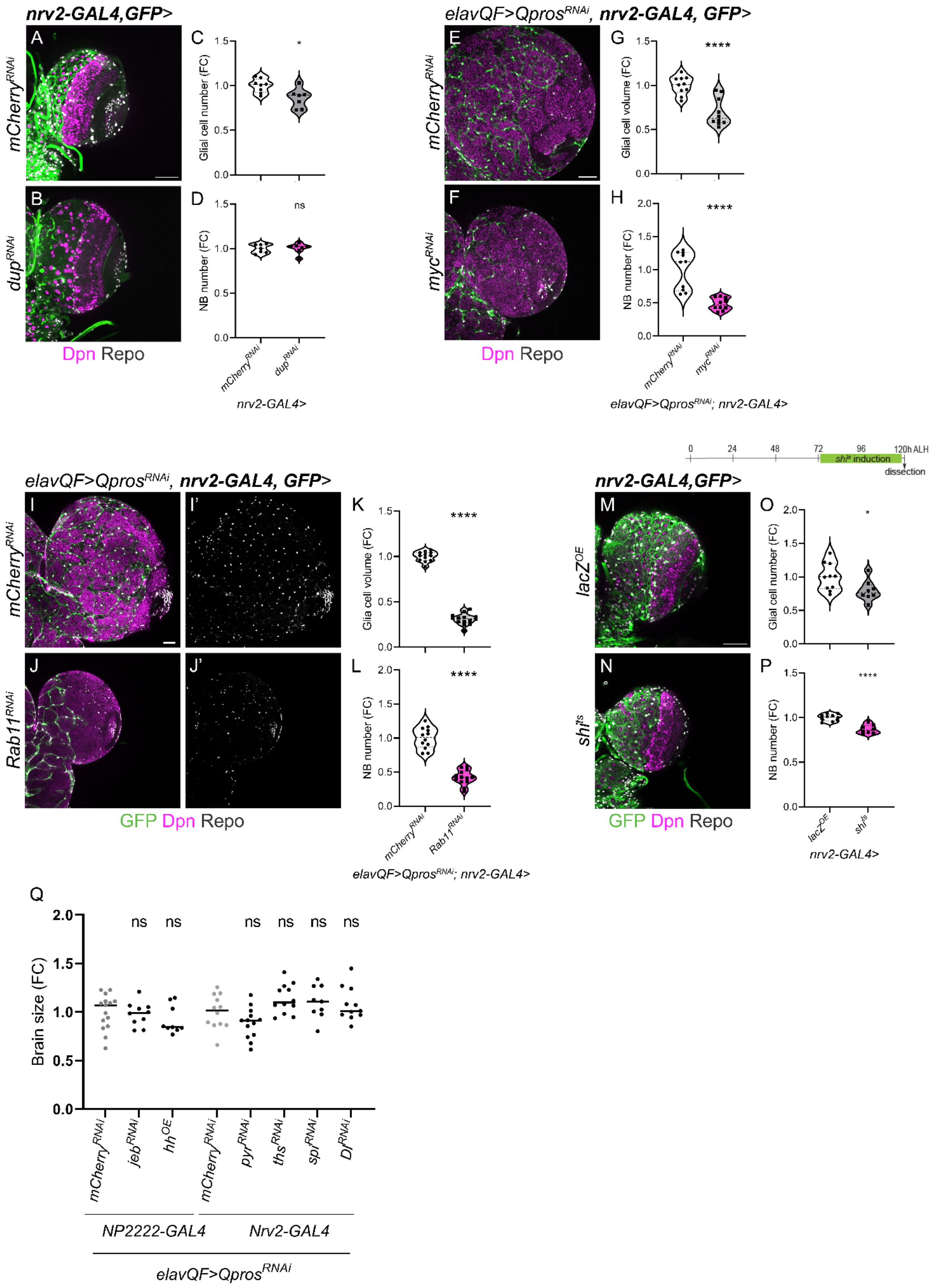
nrv2-GAL4,GFP>
elavQF>QprosRNAi, nrv2-GAL4, GFP>
Dpn Repo
nrv2-GAL4>
elavQF>QprosRNAi; nrv2-GAL4>
shits induction
dissection
120h ALH
nrv2-GAL4,GFP>
GFP Dpn Repo
Glial cell number (FC)
NB number (FC)
Glial cell volume (FC)
Glia cell volume (FC)
Brain size (FC)
NP2222-GAL4
Nrv2-GAL4
elavQF>QprosRNAi

**Figure S5. Disrupting the CG niche limits NB tumour growth.**

A-B. Representative images of brain lobes where *dup*$^{RNAi}$ was expressed in CG cells using *nrv2-GAL4, UAS-GFP*, causing a disruption of the glial niche (membranes marked in GFP and nuclei in Repo) but no effects on NB number (Dpn). Scale bar= 50 µm.

C. Quantification of glial cell number (Repo$^+$, FC) in A-B, performed using an unpaired t-test. Data normalised to controls. *mCherry*$^{RNAi}$: n= 8, m=1 ± 0.03. *dup*$^{RNAi}$: n= 7, m= 0.85 ± 0.04. *$p < 0.05$.

D. Quantification of the NB number (Dpn$^+$, FC) in A-B, performed using an unpaired t-test. Data normalised to controls. *mCherry*$^{RNAi}$: n= 7, m=1 ± 0.02. *dup*$^{RNAi}$: n= 10, m= 1 ± 0.02. ns (not significant) $p > 0.05$

E-F. Representative images of brain lobes where *myc*$^{RNAi}$ was expressed in CG cells using *nrv2-GAL4, GFP* in *elavQF>pros*$^{RNAi}$, causing a reduction in tumour size. NBs are marked with Dpn, glial membranes with GFP and nuclei with Repo. Scale bar= 50 µm.

G. Quantification of glial cell volume (Repo$^+$, FC) in E-F, performed using an unpaired t-test. Data normalised to controls. *mCherry*$^{RNAi}$: n= 10, m=1 ± 0.03. *myc*$^{RNAi}$: n= 10, m= 0.7 ± 0.05. ****$p < 0.0001$.

H. Quantification of NB number (Dpn$^+$, FC) in E-F, performed using a Mann-Whitney test. Data normalised to controls. *mCherry*$^{RNAi}$: n= 10, m=1 ± 0.09. *myc*$^{RNAi}$: n= 10, m= 0.49 ± 0.03. ****$p < 0.0001$.

I-J'. Representative images of brain lobes showing that the expression of *Rab11*$^{RNAi}$ in CG cells using *nrv2-GAL4, UAS-GFP* in *elavQF>pros*$^{RNAi}$ brains causes a reduction in tumour size and glial cell number. NBs are marked with Dpn, glial nuclei with Repo, and glial membranes with GFP. Scale bar= 50 µm.

K. Quantification of glial cell volume (Repo$^+$, FC) in I-J', performed using an unpaired t-test. Data normalised to controls. *mCherry*$^{RNAi}$: n= 12, m=1 ± 0.02. *Rab11*$^{RNAi}$: n= 14, m= 0.31 ± 0.02. ****$p < 0.0001$.

L. Quantification of NB number (Dpn$^+$, FC) in I-J', performed using an unpaired t-test. Data normalised to controls. *mCherry*$^{RNAi}$: n= 12, m=1 ± 0.02. *Rab11*$^{RNAi}$: n= 14, m= 0.31 ± 0.02. ****$p < 0.0001$.

M-N. Representative images of brain lobes where *shi*$^{ts}$ was expressed in CG cells from 72 h AEL using *nrv2-GAL4, GFP*, causing a reduction in glial (Repo) and NB number (Dpn), compared to control (*lacZ*$^{OE}$). Scale bar= 50 µm.

O. Quantification of glial cell number (Repo$^+$, FC) in M-N, performed using an unpaired t-test. Data normalised to controls. *lacZ*$^{OE}$: n= 10, m=1 ± 0.07. *shi*$^{ts}$: n= 8, m= 0.8 ± 0.05. *$p < 0.05$.

P. Quantification of the NB number (Dpn$^+$, FC) in M-N, performed using an unpaired t-test. Data normalised to controls. *lacZ*$^{OE}$: n= 10, m=1 ± 0.01. *shi*$^{ts}$: n= 8, m= 0.87 ± 0.02. ****$p < 0.0001$.

Q. Knockdown of candidate secreted factors in CG cells using *NP2222-GAL4 or nrv2-GAL4* in *elavQF>Qpros*$^{RNAi}$ tumours resulted in no significant reduction in brain tumour size. Data normalised to controls. Analysis for *NP2222-GAL4* data was performed using a Kruskal-Wallis test. *mCherry*$^{RNAi}$: n= 15, m=1 ± 0.05. *jeb*$^{RNAi}$: n= 10, m= 0.98 ± 0.04. *hh*$^{OE}$: n= 9, m= 0.92 ± 0.05. Analysis for *Nrv2-GAL4* data was performed using an ordinary one-way ANOVA. *mCherry*$^{RNAi}$: n= 12, m=1 ± 0.05. *pyr*$^{RNAi}$: n= 13, m= 0.88 ± 0.04. *ths*$^{RNAi}$: n= 12, m=1.14 ± 0.04. *spi*$^{RNAi}$: n= 9, m= 1.1 ± 0.06. *Dl*$^{RNAi}$: n= 11, m=1.08 ± 0.05. ns (not significant) $p > 0.05$.

**Table 1. Fly strains and sources.**

| Strain | Source | Stock centre/ Reference/Comments |
|---|---|---|
| *$w^{1118}$* | BDSC | |
| *elav-QF2* | BDSC | 66466 |
| *dnab-GAL4* | Gould lab | |
| *Insc-GAL4, wor-GAL4* | Gould lab | |
| *repo-GAL4, tubGAL80$^{ts}$* | BDSC | 7415 |
| *NP2222-GAL4* | Kyoto DGGR | 112830 |
| *Nrv2-GAL4, UAS-GFP* | BDSC | 6795 |
| *wrapper-GAL4* | Marshall lab | |
| *GMR85G01-GAL4* | BDSC | 40436 |
| *Poxn-GAL4, UAS-GFP* | Maurange lab | Narbonne et al |
| *tubGAL80$^{ts}$* | BDSC | |
| *tub-Gal4, UAS-nlsGFP::6xmyc::NLS, hs-flp; FRT82B, tubP-Gal80 LL3/TM6B* | | Lee and Luo, 1999 |
| *FRT82B, pros$^{17}$* | BDSC | |
| *UAS-RFP.nls* | BDSC | 30556 |
| *UAS-GFP.nls* | | |
| *gcm-lacZ* | BDSC | 5445 |
| *G-TRACE* | BDSC | 28280 |
| *UAS-FUCCI* | BDSC | 55114 |
| *UAS-lacZ* | BDSC | 8529 |
| *UAS-lacZ$^{RNAi}$* | BDSC | |
| *UAS-mCherry$^{RNAi}$* | BDSC | 35785 |
| *UAS-Dcr-2* | BDSC | 24651 |
| *UAS-pros$^{RNAi}$* | BDSC | 42538 |
| *QUAS-pros$^{RNAi}$* | This study | |
| *UAS-aPKC$^{CAAX}$* | Richardson Lab | |
| *UAS-brat$^{RNAi}$* | BDSC | 34646 |
| *UAS-repo* | | |
| *UAS-repo$^{RNAi}$* * | BDSC | 28339 |
| *UAS-gcm$^{DN}$* | Giangrande Lab | |
| *UAS-hid, rpr* | G. Viktorin | |
| *UAS-shi$^{ts}$* | BDSC | 44222 |
| *UAS-rab11$^{RNAi}$* | BDSC | 27730 |
| *UAS-sec6$^{RNAi}$* | VDRC | 22077 |
| *UAS-htl$^{DN}$* | BDSC | 5366 |
| *UAS-dsor$^{RNAi}$* | VDRC | 40026 |
| *UAS-PTEN* | Xu lab | |
| *UAS-syp* | Tzumin Lee Lab | |
| *UAS-syp$^{RNAi}$* | BDSC | 56972 |
| *UAS-EcR$^{DN\ F645A}$* | Quinn lab | |
| *UAS-myc$^{RNAi}$* | VDRC | 106066 |

| | | |
|---|---|---|
| *UAS-Dap* | Lehner lab | |
| *UAS-Rbf* | BDSC | 50748 |
| *UAS-dup*$^{RNAi}$ | BDSC | 67210 |
| *UAS-jeb*$^{RNAi}$ | BDSC | 56022 |
| *UAS-hh*$^{OE}$ | Kornberg lab | |
| *UAS-pyr*$^{RNAi}$ | VDRC | 36523 |
| *UAS-ths*$^{RNAi}$ | VDRC | 24536 |
| *UAS-spi*$^{RNAi}$ | VDRC | 3922 |
| *UAS-Dl*$^{RNAi}$ | BDSC | 36784 |

* Tested with *repo-GAL4* - caused embryonic lethality

**Table 2. Mean, SEM, sample size (n) and adjusted p-value for Figure S1 A.**

| | *elavQF>+* | | | *elavQF>Qpros*$^{RNAi}$ | | | **p-value** |
|---|---|---|---|---|---|---|---|
| **h AEL** | **Mean** | **SEM** | **n** | **Mean** | **SEM** | **n** | |
| 0 | 0 | 0 | 4 | 0 | 0 | 4 | ns >0.9999 |
| 24 | 0 | 0 | 4 | 0 | 0 | 4 | ns >0.9999 |
| 48 | 0 | 0 | 4 | 0 | 0 | 4 | ns >0.9999 |
| 72 | 0 | 0 | 4 | 0 | 0 | 4 | ns >0.9999 |
| 96 | 0 | 0 | 4 | 0 | 0 | 4 | ns >0.9999 |
| 120 | 42.5 | 7.5 | 4 | 0 | 0 | 4 | ****<0.0001 |
| 144 | 100 | 0 | 4 | 2.5 | 1.60 | 4 | ****<0.0001 |
| 168 | 100 | 0 | 4 | 11.67 | 3.97 | 4 | ****<0.0001 |
| 192 | 100 | 0 | 4 | 48.33 | 6.45 | 4 | ****<0.0001 |
| 216 | 100 | 0 | 4 | 69.17 | 6.85 | 4 | ****<0.0001 |

**Table 3. Mean, SEM and sample size (n) for Figure 2 J.**

| | **Mean** | **SEM** | **n** | **p-value** | **Mean** | **SEM** | **n** | **p-value** | **Mean** | **SEM** | **n** | **p-value** |
|---|---|---|---|---|---|---|---|---|---|---|---|---|
| **72 h AEL** | G1 | | | | S | | | | G2/M | | | |
| Control | 9.67 | 1.68 | 10 | ns = 0.854 | 50.46 | 6.07 | 10 | ns = 0.181 | 39.88 | 5.06 | 10 | ns = 0.574 |
| *elavQF> pros*$^{RNAi}$ | 14.34 | 2.35 | 8 | | 38.22 | 5.55 | 8 | | 47.44 | 4.22 | 8 | |
| **96 h AEL** | G1 | | | | S | | | | G2/M | | | |
| Control | 7.86 | 1.61 | 10 | ns = 0.655 | 52.45 | 5.95 | 10 | ns = 0.226 | 39.68 | 5.20 | 10 | ns = 0.651 |
| *elavQF> pros*$^{RNAi}$ | 13.82 | 2.52 | 9 | | 40.52 | 4.03 | 9 | | 45.67 | 1.96 | 9 | |
| **120 h AEL** | G1 | | | | S | | | | G2/M | | | |
| Control | 95.68 | 0.52 | 13 | **** <0.0001 | 1.58 | 0.36 | 13 | **** <0.0001 | 2.74 | 0.45 | 13 | **** <0.0001 |
| *elavQF> pros*$^{RNAi}$ | 28.26 | 3.54 | 10 | | 14.14 | 3.43 | 10 | | 57.60 | 2.19 | 10 | |

**Table 4. Mean, SEM and sample size (n) for Figure 2 O.**

| | Mean | SEM | n | p-value | Mean | SEM | n | p-value | Mean | SEM | n | p-value |
|---|---|---|---|---|---|---|---|---|---|---|---|---|
| | **72 h AEL** | | | | **96 h AEL** | | | | **120 h AEL** | | | |
| Control | 97.3 | 5.63 | 10 | ** = 0.0026 | 273.1 | 10.43 | 10 | **** <0.0001 | 331.2 | 21.92 | 13 | **** <0.0001 |
| *elavQF> pros*$^{RNAi}$ | 130.5 | 6.81 | 8 | | 457.7 | 23.21 | 9 | | 1034 | 38.71 | 10 | |

**Table 5. Mean, SEM, sample size (n) and adjusted p-value for Figure 3 F.**

| | *elavQF>+* | | | *elavQF>Qpros*$^{RNAi}$ | | | p-value |
|---|---|---|---|---|---|---|---|
| **h AEL** | **Mean** | **SEM** | **n** | **Mean** | **SEM** | **n** | |
| 48 | 0 | 0 | 4 | 0 | 0 | 4 | ns >0.9999 |
| 72 | 0 | 0 | 4 | 0 | 0 | 4 | ns >0.9999 |
| 96 | 0 | 0 | 4 | 0 | 0 | 4 | ns >0.9999 |
| 120 | 0 | 0 | 4 | 43.61 | 4.72 | 4 | ****<0.0001 |
| 144 | 0 | 0 | 4 | 79.72 | 6.94 | 4 | ****<0.0001 |
| 164 | 22.5 | 8.54 | 4 | 97.22 | 2.78 | 4 | ****<0.0001 |
| 188 | 71.43 | 13.35 | 4 | 97.22 | 2.78 | 4 | **=0.0041 |

**Table 6. Mean, SEM and sample size (n) for Figure 3 I.**

| | Mean | SEM | n | p-value | Mean | SEM | n | p-value | Mean | SEM | n | p-value |
|---|---|---|---|---|---|---|---|---|---|---|---|---|
| **120 h AEL** | G1 | | | | S | | | | G2/M | | | |
| Ethanol | 39.7 | 5.38 | 11 | **** <0.0001 | 16.89 | 2.80 | 11 | * = 0.0297 | 43.42 | 5.22 | 11 | **** <0.0001 |
| 20E | 95.24 | 1.4 | 7 | | 0.97 | 0.4 | 7 | | 3.78 | 1.48 | 7 | |

**Table 7. Mean, SEM and sample size (n) for Figure 3 L.**

| | Mean | SEM | n | p-value | Mean | SEM | n | p-value | Mean | SEM | n | p-value |
|---|---|---|---|---|---|---|---|---|---|---|---|---|
| **120 h AEL** | G1 | | | | S | | | | G2/M | | | |
| *lacZ*$^{RNAi}$ | 68.1 | 8.38 | 6 | **** <0.0001 | 2.47 | 1.45 | 6 | * = 0.0291 | 29.42 | 9.19 | 6 | * = 0.0163 |
| *EcR*$^{DN}$ | 21.81 | 3.05 | 6 | | 24.67 | 4.55 | 6 | | 53.51 | 2.74 | 6 | |

**Table 8. Mean, SEM and sample size (n) for Figure 3 O.**

| | Mean | SEM | n | p-value | Mean | SEM | n | p-value | Mean | SEM | n | p-value |
|---|---|---|---|---|---|---|---|---|---|---|---|---|
| **144-168 h AEL** | G1 | | | | S | | | | G2/M | | | |
| *lacZ*$^{RNAi}$ | 90.26 | 0.96 | 12 | **** <0.0001 | 1.83 | 0.41 | 12 | ***= 0.0004 | 7.91 | 0.96 | 12 | **** <0.0001 |
| *EcR*$^{DN}$ | 51.94 | 3.63 | 12 | | 13.82 | 2.11 | 12 | | 34.24 | 2.61 | 12 | |

**Table 9. Primers used in this study.**

| Name | Sequence (5'-3') |
|---|---|
| **Transgenic *QUAS-pros*$^{RNAi}$ line** | |
| pros BL42538 top | CTAGCAGTCAGGATGTGGAACAAGAACAATAGTTATATTCA AGCATATTGTTCTTGTTCCACATCCTGGCG |
| pros BL42538 bottom | AATTCGCCAGGATGTGGAACAAGAACAATATGCTTGAATAT AACTATTGTTCTTGTTCCACATCCTGACTG |
| **RT-qPCT** | |
| Rpl32 FOR | CCGCTTCAAGGGACAGTATCTG |
| Rpl32 REV | ATCTCGCCQCAGTAAACGC |
| Phm FOR | GGCATCATGGGTGGATTT |
| Phm REV | CAAGGCCTTTAGCCAATCG |
| Spok FOR | GCGGTGATCGAAACAACTC |
| Spok REV | CGAGCTAAATTTCTCCGCTTT |
